# Wide-field time-gated SPAD imager for phasor-based FLIM applications

**DOI:** 10.1101/687277

**Authors:** Arin Ulku, Andrei Ardelean, Michel Antolovic, Shimon Weiss, Edoardo Charbon, Claudio Bruschini, Xavier Michalet

**Affiliations:** AQUA Lab, Ecole Polytechnique Fédérale de Lausanne, Neuchâtel, Switzerland; Department of Chemistry & Biochemistry, UCLA, Los Angeles, CA, USA

## Abstract

We describe the performance of a new wide area time-gated single-photon avalanche diode (SPAD) array for phasor-FLIM, exploring the effect of gate length, gate number and signal intensity on the measured lifetime accuracy and precision. We conclude that the detector functions essentially as an ideal shot noise limited sensor and is capable of video rate FLIM measurement. The phasor approach used in this work appears ideally suited to handle the large amount of data generated by this type of very large sensor (512×512 pixels), even in the case of small number of gates and limited photon budget.

## 1. Introduction

Among the many optical imaging modalities available today, fluorescence imaging is particularly popular in biological sciences due to its versatility and specificity. Fluorescence imaging can indeed target almost any molecule of interest with minimal interference with the molecule’s function, while a vast range of fluorophores with distinct absorption and emission spectra is available today. This has made it the tool of choice for multiplexed imaging, with applications ranging from DNA sequencing, diagnostics, cell imaging, superresolution microscopy, and especially *in vivo* imaging for longitudinal (pre-) clinical studies of diseases and therapy monitoring.

Fluorescence lifetime imaging adds an important dimension to conventional fluorescence microscopy, by directly measuring the de-excitation rate of the fluorophore, in addition to its intensity (1-3). The lifetime is the inverse of that rate, which comprises not only the radiative rate (corresponding to the emission of a photon), but also all the non-radiative de-excitation rates. This makes the fluorescence lifetime sensitive to changes in any of these non-radiative rates, some of which are dependent on the fluorophore’s electronic environment. A particularly attractive property of this measurement is that it is direct and in most circumstances, independent on the signal level, thus making it an extremely sensitive molecular probe of the local environment.

Fluorescence lifetime can be measured in many different ways, using for instance frequency modulation, time-correlated single-photon counting (TCSPC) or time-gating, in combination with pulsed laser excitation (4). Most commercial devices offering fluorescence lifetime imaging microscopy (FLIM) are single-spot beam scanning confocal microscopes, which providing 3-dimensional sectioning. However, relatively long acquisition times and high excitation powers are generally needed. Both are detrimental to live imaging, as raster scanning with long acquisition time results in the possibility that the sample moves between the acquisition start and end, and high excitation powers can result in premature photobleaching and photodamage of the sample.

Wide-field FLIM techniques, which acquire data from every points in the field of view simultaneously, solve some of the throughput and photodamage issues, although they come with challenges of their own (5). The most established of these technologies uses time-gated intensified CCD (or CMOS) cameras (ICCD/ICMOS), scanning the fluorescence decay in the time domain to acquire data from “time slices” covering the whole laser period (6). Technologically similar, but based on a fundamentally different principle, position-sensitive, photon-counting detectors allow TCSPC measurements to be performed (7-9). Finally, single-photon avalanche diode (SPAD) arrays, notably those designed using standard CMOS processes (discussed in more detail below), can be found as either TCSPC or time-gated variants, the latter being more common for very large arrays, due to the challenge of implementing massive TCSPC electronics on the chip itself (e.g. (10-12), reviewed in (13)).

No matter which technique is used, FLIM data poses additional challenges, such as data storage, processing and representation. Instead of a mere intensity value per pixel, FLIM data is either comprised of list of photon time stamps (TCSPC approaches) or of many binned or time-gated intensity values (TCSPC approaches resulting in binned data, or time-gated approaches), from which one or more lifetimes and their corresponding amplitudes needs to be extracted and represented. Over the years, many different data analysis and representation approaches have been proposed, most of which rely on the assumption that each fluorescent species can be described by a single-exponential decay (14,15). While this is appropriate in many cases, it cannot be generalized, and a more model-agnostic approach such as phasor analysis (16-19) has many advantages. Phasors are easily calculated and their graphical representation allows simple interpretation and localization of species with different decays, while not precluding accurate quantitative analysis.

Herein, we describe applications of a very large time-gated CMOS SPAD camera for phasor-FLIM, SwissSPAD2(20), and discuss the main parameters affecting its performance. We conclude with a brief overview of future prospects for the technology.

## 2. Experimental/Methods

### 2.1. Technical Overview of SwissSPAD2

The detector used in this article is SwissSPAD2 (SS2), a high-speed, large-format SPAD imaging sensor with a time-gate integrated on the same chip (20). The sensor chip consists of 512×512 pixels, of which only 472×256 pixels were enabled in the camera module tested here. The pixel pitch is 16.38 μm, and the crosstalk probability between neighboring pixels is less than 0.075% (Fig. S4). Because of the digital nature of each pixel (a photon is detected, or none), the camera captures binary images with ideally no readout noise, making it suitable for single-photon imaging. Each pixel has a 1-bit memory electronics, the whole array being read at a maximum speed of 97.7 kfps (kilo frames per second). Each sequence of 255 binary frames is accumulated into an 8 bit gate image on field-programmable gate-array (FPGA), and transferred to the data acquisition memory of a PC via a USB 3.0 connection. More detailed technical specifications of SS2 were reported in (20).

SS2 performs time-resolved imaging using its in-pixel gate electronics. The global (array-wide) gate signal is generated, using mixed-mode clock manager (MMCM) modules on FPGA, from the laser trigger signal transmitted to the camera by the laser controller (or a fast laser pick-up PIN diode). Briefly, during each 1-bit frame exposure (user-selectable in multiple of 400 ns, minus 50 ns), the gate is turned on and off after each laser pulse, and any detected photon sets the pixel memory to 1. If more than one photon is detected, the subsequent photons are ignored. After the set exposure time, the 1-bit frame is readout, and the procedure is repeated until the user-defined total number of frames has been acquired (typically 255 – for an 8 bit gate image, or 4×255 for a 10 bit gate image). The accumulated gate image is then transferred to the PC, while a new gate position is defined and the procedure repeated to acquire a new gate image, and so on, until the requested number of gate images has been acquired.

SS2’s gate duration *W* is significantly longer (> 10 ns) than most common fluorophore lifetimes, but can be triggered very precisely with respect to the laser pulse, using steps of 17.9 ps. Fig. 1 illustrates the characteristics of a typical gate window. This gate profile was measured by recording the detector’s response to a 20 MHz pulsed laser using gate images stepped by 17.9 ps through the 50 ns laser period. The figure shows a window spanning 70 ns, but the gate profile is periodic with a period of 50 ns.

**Fig. 1.**
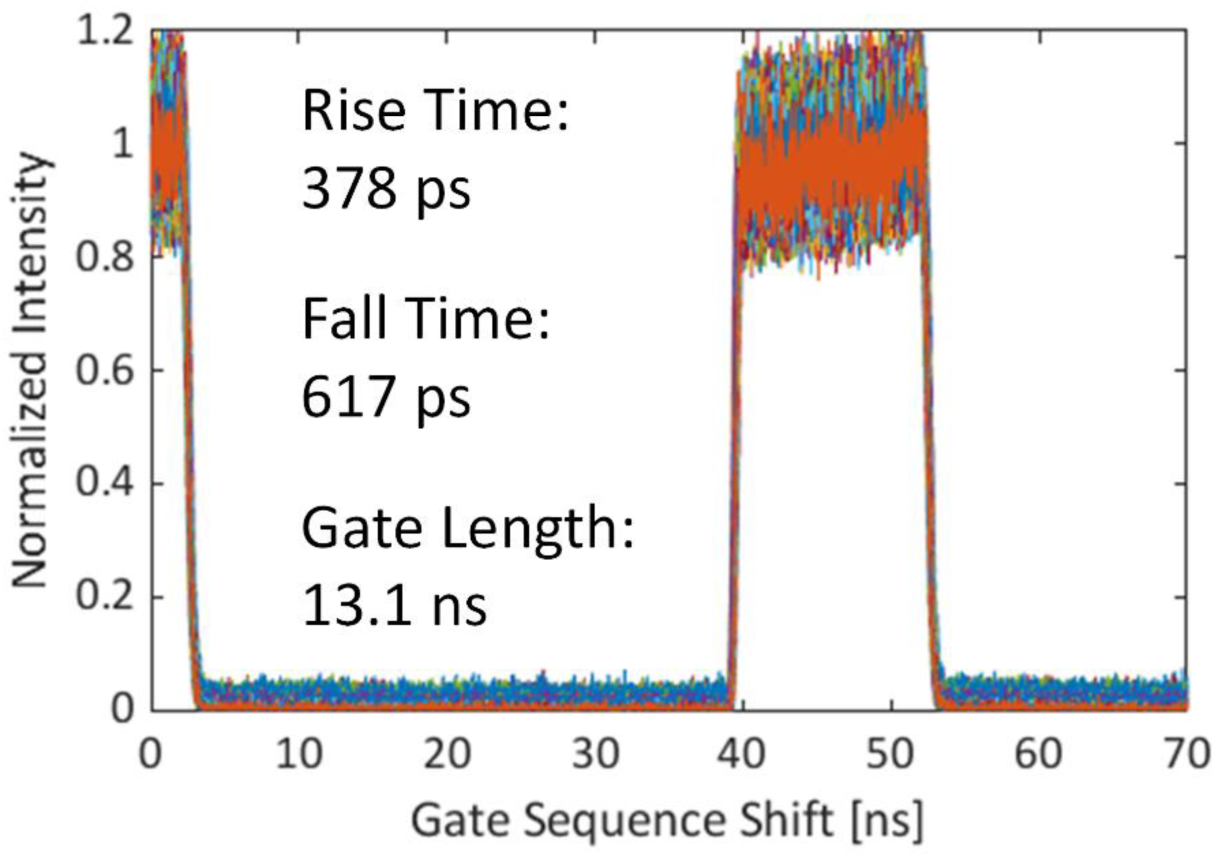
Characteristics of the gate in the firmware that was used in the FLIM experiment. The response of every other 4th pixel in the center 472×256 array is plotted. The minimum achievable gate length is 10.8 ns.

Seven gate configurations whose gate length *W* was varied between 10.8 ns and 22.8 ns were tested in these experiments. The gate length and position determine the temporal window during which the SPAD is sensitive after each laser pulse. At fixed laser frequency and intensity, a wider gate can collect more photons over a given exposure time, at the expense of a lower photon arrival time resolution. As we shall see, this is not a fundamental limit. The software allows selecting the gate configuration (length), the number of laser pulses per 1-bit frame (exposure), the bit depth (8 or 10 bits) of each gate image (dynamic range), the delay between two successive gate positions (step), and the number of gate images in the dataset. The characteristics of the gate configurations tested here are listed in Table S1.

The gate properties affect the time-resolved imaging performance and can influence the fluorescence life-time determination accuracy and precision. For wide-field systems, the spatial uniformity of the measurements is determined by the gate edge position distribution, or skew. In large-format sensors, the skew of the gate signals and possible voltage drops during high-frequency signal toggling cause gate edge non-uniformity across the array (20). As the gate length increases, a significant narrowing of the rising edge skew is observed (next to last line in Table S1). This effect can be attributed to the difference in the supply voltage fluctuation level during signal transitions. The first gate signal transition (which corresponds to the falling edge of the gate window since the gate moves forward with respect to the laser trigger) causes a spatially uneven supply voltage drop in the gate signal trees, which results in a skew in the second gate signal transition, in this case the rising edge. As the gate length increases, the better recovery in the voltage drop over a longer delay between transitions reduces the skew. Since this source of gate non-uniformity is deterministic, it can be corrected by calibration after the measurements, as described in the next section.

Two other key parameters of the gate performance are the rise and fall times. Their main contributors are laser pulse width, SPAD response and gate signal jitter, as well as the switching speed of the gate transistor. The latter is determined by fabrication process constraints. The steepness of the gate edge also depends on the readout speed and the laser frequency due to the variation of the supply voltage swing with these parameters. The timing resolution is therefore affected by a series of stochastic effects over some of which we do not have control, and therefore their influence was not studied in this work.

The 10.5% native fill factor of SwissSPAD2 can be partially compensated for by microlenses. Fig. 2a shows a microscope image of a SwissSPAD2 array with a microlenses deposited on the pixels using a procedure described in (21). The microlens performance at normal angle of incidence was measured in a fluorescence microscope (IX81, Olympus, Japan). In this experiment, the image of a *convallaria majalis* sample was captured successively with two SwissSPAD2 cameras (one with and one without microlenses), using the same camera exposure and illumination settings. The concentration factor (CF), defined as the ratio *μ*_*m*_/*μ*_*nm*_, where *μ*_*m*_ and *μ*_*nm*_ are the mean photon counts of the camera with and without microlenses after subtraction of the detector dark counts, was found to be CF = 2.65, corresponding to an effective fill factor of 27.8.

**Fig. 2.**
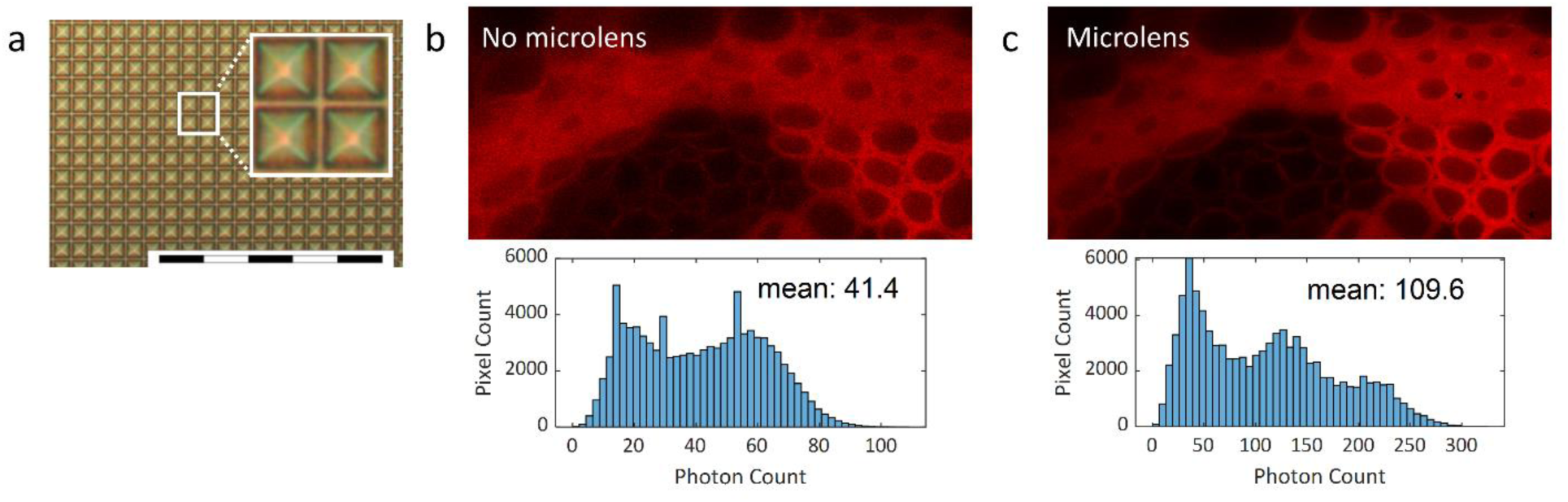
(a) Microscopic image of SwissSPAD2 pixels with microlenses. Scale bar is 200 μm. (b-c) Fluorescence intensity image of a *convallaria majalis* sample captured with SwissSPAD2 (b) without and (c) with microlenses (22). Mean photon count without microlenses: 41.4. Mean photon count with microlenses: 109.6. Microlens concentration factor: 2.65. Experimental parameters: V_ex_: 6.5 V, array size: 453×210, bit depth: 10, integration time: 3.21 ms, *λ*_emission_: 607 nm, pile-up correction: on. Hot pixels with 1% highest dark count rate in the array were corrected using an interpolation method based on setting their intensity values to the mean of the four nearest-neighbor pixels.

As this concentration factor is lower than the theoretically calculated value (21), we tested the two sensors on a simple optical setup where the angle between the sensor and a collimated laser beam (785 nm, PiLas, A.L.S., Germany) can be adjusted in both dimensions. Both sensors were tested successively, and the total photon counts measured as a function of incidence angle. The maximum concentration factor, calculated in this way, was found to be 4.46 at the optimal angle at 3.5 V excess bias voltage. This difference with the normal incidence’s CF could be due to a slight misalignment of the microlens array with respect to the SPAD array or local variations in microlens characteristics (21).

Table 1 summarizes the performance of SS2 and compares it with other state-of-the-art, large-format scientific cameras. SPAD cameras have ideally zero readout noise as a result of their digital nature; therefore they can perform wide-field FLIM with single-photon sensitivity. Their CMOS technology is scalable, robust and cost-effective compared to MCPs and photocathode-based detectors. Among SPAD cameras, SS2 employs the largest array size to date, which enables both wide field of view and high spatial resolution.

**TABLE 1:** Comparison of SwissSPAD2’s performance with state-of-the-art time-resolved scientific cameras. Some detectors are (or can be) equipped with microlenses (µL) and have therefore two distinct fill factor values: native (without microlenses) and with microlenses. The Maximum PDP/QE value correspond to the case with microlenses when applicable.

| Source | SwissSPAD2<br>(This Work &<br>(22)) | Ref. (36) | Ref. (37) | Ref. (38) | Ref. (12) | SS1 (11) | H33D Gen I<br>(9) | LaVision<br>PicoStar HR (39) | Ref. (40) |
| --- | --- | --- | --- | --- | --- | --- | --- | --- | --- |
| **Detector Type** | SPAD | SPAD | SPAD | SPAD | SPAD | SPAD | MCP | Gated ICCD | CMOS Image<br>Sensor |
| **Array Format** | 512×512<br>472×256 used | 320×240 | 160×120 | 256×256 | 256×256 | 512×128 | Ø 400 | 1,376×1,040 | 256×512 |
| **Pixel Size/Spatial**<br>**Resolution** | 16.38 $\mu\text{m}$ | 8 $\mu\text{m}$ | 15 $\mu\text{m}$ | 8 $\mu\text{m}$ | 16 $\mu\text{m}$ | 24 $\mu\text{m}$ | 100 $\mu\text{m}$ | 6.45 $\mu\text{m}$ | 11.2 $\mu\text{m}$ (H)<br>× 5.6 $\mu\text{m}$ (V) |
| **Fill Factor**<br>**(Native)** | 10.5% | 26.8% | 21% | 19.6% | 61% | 5% | ~100% | ~100% | 16.7% |
| **Fill Factor**<br>**(with $\mu\text{L}$)** | >28-47% | 50% | - | - | - | 60% | - | - | - |
| **rms Readout**<br>**Noise** | 0 | 0.168 $\text{e}^-$ | - | - | 0 | 0 | 0 | 4-6 $\text{e}^-$ | 1.75 $\text{e}^-$ |
| **Maximum**<br>**PDP/QE** | ~50%<br>@520 nm | 39.5%<br>@480 nm | - | - | 39.5% @480<br>nm | 46%<br>@490 nm | 18%<br>@400 nm<br>(S20) | 11%<br>@550 nm<br>(S25) | - |
| **Minimum Gate**<br>**Length** | 10.8 ns | - | 10 ns | - | 4 ns | 4 ns | - | 200 ps | 12.7 ns |
| **Minimum Gate**<br>**Delay Step** | 17.9 ps | - | 194 ps | - | - | 20 ps | 150 ps | 5 ps | 1 ns |
| **Dark Count**<br>**(median)** | 7.5 cps/px<br>@27°C | 47 cps/px | 580 cps/px | 50 cps/px | 6.2 kcps/px | 366 cps/px | 20 kHz<br>(global) | < 0.1 $\text{e}^-/\text{px}/\text{s}$<br>(CCD) | - |
| **Maximum Frame**<br>**Rate** | 97.7 kfps<br>(1 bit) | 16 kfps<br>(1 bit) | 486 fps<br>(5.4 bit) | 4 kfps<br>(3 bin<br>histogram) | 100 kfps<br>(1 bit) | 156 kfps<br>(1 bit) | Shot noise-limited,<br>700 kcps | 10 fps<br>(12 bit) | 12 fps |

### 2.2. Data pre-processing

#### Pile-up correction

As a result of the 1-bit storage scheme of the camera, the recorded signal does not scale linearly with the incident signal. Instead, the percentage of unrecorded photons increases with incident signal, a phenomenon known as pile-up. This leads to an artificial non-linearity (saturation) of the recorded signal, disproportionally affecting gates with high count values compared to gates with lower counts. The net result is a deformation of the recorded decay shape compared to its expected shape, ultimately causing an error in the calculated lifetime unless corrected for. This effect is easily accounted for by the following correction formula (22):

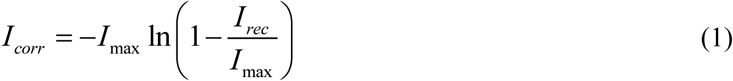

where *I*_*corr*_ is the corrected photon count, *I*_*rec*_ is the recorded photon count and *I*_*max*_ is the maximum photon count that can be recorded, equal to the number of binary frames in the gate image (255 for a 8-bit image or 255×4 = 1,020 for a 10-bit image). While this correction method is useful to recover the incident decay profile, pile-up also decreases the signal-to-noise ratio (SNR) in a non-trivial manner. Assuming a standard Poisson-distributed incident signal, the SNR calculated from the pile-up corrected signal, 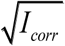, the closest estimation we can compute, is ignoring the departure from a Poisson distribution resulting from the pile-up process, and underestimates the noise level resulting from it. For simplicity, we do not consider this effect in our analysis.

#### Background correction

Uncorrelated background signal due to detector noise must be taken into account when computing phasors. There are multiple effects of uncorrelated background. First, it affects the phasor location in a non-trivial manner; it also degrades the precision of the results by increasing the standard deviation of the photon count; finally, it contributes to signal saturation due to pile-up. In this work, uncorrelated background was estimated and subtracted from the pile-up corrected signal as described in (23). Briefly, the average value of gates covering a region of the laser period where the fluorescence contribution is negligible (the tail of the decay in our experiments), was used for a sample whose lifetime was significantly shorter than *T – W*. For decay characterized by longer lifetimes such as the samples used here, the decay never reaches the background level due to the periodic excitation, and a different approach based on a 3-point background estimation needs to be used (24). While these methods do not mitigate the slight increase in dispersion caused by background photons, they improve the calculated phasor location and subsequent analysis.

#### Noisy pixel consideration

Due to imperfection in the fabrication process, a small percentage of the SPADs in the array are characterized by high dark count rates (> 1,000 counts/s or 1 kcps). The pile-up and background corrections described above should in principle take care of the resulting offset, but, due to saturation, this correction may increase the resulting uncertainty on the signal. To eliminate their influence on phasor calculation (and its variance), pixels with values in the top intensity percentile can be rejected if desired. However, this procedure also rejects naturally brighter regions of the sample and is therefore not always recommended. We found that single pixel phasor values corresponding to such noisy pixels are easily distinguished from the remainder of the pixels as they are randomly scattered in the phasor plot (generally on the lien connecting sample fluorescence and uncorrelated background). For analyses using binned data (region of interest or ROI) rather than single pixel values, their influence on the calculated ROI phasors is reduced and therefore was ignored in most cases.

### 2.3. Frame rate definition

In raw data files provided in the accompanying data repository (25), each 10-bit gate image is in fact comprised of 4 consecutive 8-bit gate images. In the global shutter mode used in these measurements, exposure and readout are performed sequentially for each binary image, and the frame rate of a sequence is calculated as the inverse of the acquisition time (sum of exposure and readout), defined as:

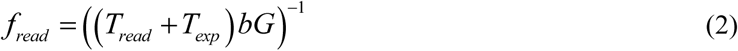

where *T*_*read*_ is the binary frame readout time, *T*_*exp*_ is the binary frame exposure time, *b* is the number of binary images in a gate position, and *G* is the number of acquired gate positions. At the highest readout speed of 472×256 pixel frames, *T*_*read*_ is equal to 10.2 μs. *b* is 255 for 8-bit gate images and 1,020 for 10-bit gate images. *T*_*exp*_ is determined by an on-FPGA counter that increments every 400 ns and has a 50 ns offset:

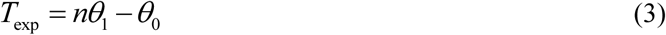

*w*here *θ*_*0*_ = 50 ns and *θ*_*1*_ = 400 ns are firmware constants and *n* is a user-selectable parameter. Since all datasets were acquired as sets of four 8-bit gate images, the actual frame rates for 8-bit gate images are four times lower than the calculated acquisition time.

As the current firmware allows continuous acquisition of at most 250 10-bit images due to USB 3.0 data transfer stability issues for long sequences, 2,800 gate positions were acquired by restarting the FPGA after each sequence of 250 gate positions. The ∼1.5 s dead time resulting from each FPGA restart introduces a further reduction of the actual frame rate, which is not reflected in the above formula. This delay would be eliminated during the acquisition of an 8-bit dataset comprising of less than 1,000 gate positions, since it would not require FPGA restart, or when the current USB transfer issues are solved.

In the experiment reported in Section 3.3, where the effect of frame rate on lifetime determination precision was analyzed (Fig. 7), the reported “virtual” frame rate was calculated using Eq. (2), but with the number *G* of gate positions retained for the analysis, rather than the number gates used to acquire the original dataset (*G* = 2,800).

### 2.4. Phasor Analysis of Time-Gated Data

#### Phasor calculation

The phasor method is a visual representation of fluorescence decay profiles on a two dimensional map, where each decay is represented by a single point *P* with coordinate (*g, s*) or equivalently, a complex number *z = g + i s* = *m e*^*iϕ*^ with unique phase *ϕ* and modulus *m*. This method was first introduced for FLIM in the frequency domain (16,26). Its use for time domain FLIM with wide-field detectors was recently illustrated with single-photon counting TCSPC detectors(19) or time-gated detectors (23,27).

In this work, we used the phasor method to analyze FLIM data obtained by SS2 with long overlapping gates. As illustrated in Fig. 3a, a gate with a width *W* (between 10.8 and 22.8 ns, depending on the measurement series) was scanned across a fluorescent decay with delay steps as short as 17.9 ps. The number *G* of gate positions, which directly influences the total acquisition time, is equal to the ratio between the laser period *T* and the gate step *δt*. Each gate position (indexed by *k*) is represented by a time stamp *t*_*k*_ = (*k - 1*) *δt*, indicating the delay between the start of the gate and the laser pulse. The phasor of the decay, *z*, is calculated according to:

**Fig. 3.**
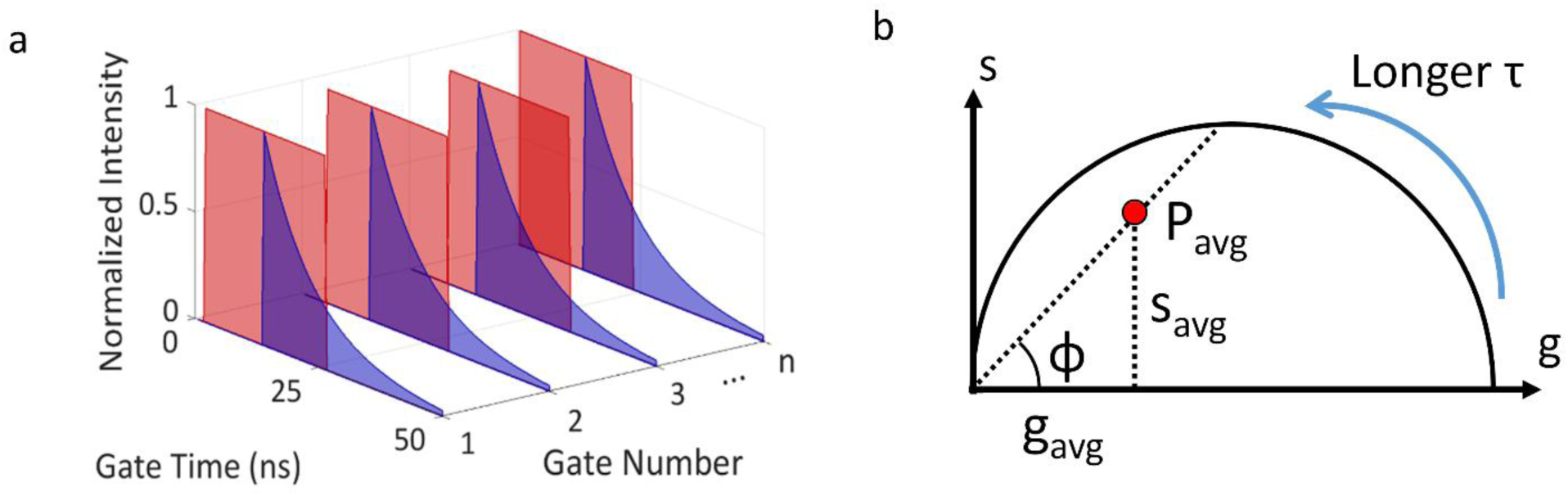
Conceptual illustration of the phasor method. (a) A gate with a fixed width W is scanned across the 50 ns fluorescence decay period. Each gate is associated with a “nanotime” specifying its start time with respect to the laser pulse. Each pixel in a gate image contains the number of photons detected during the gate image exposure time. (b) The phasor of the decay (*P*) recorded in a given pixel is calculated as the weighted average of the gate image intensity multiplied by a cosine or sine term depending on the gate nanotime (Eq. (3)). For a single-exponential decay, *P* is located on the universal semicircle, approaching the origin point (0, 0) as lifetime increases toward infinity. The phase lifetime is calculated using *ϕ*, the angle of the line connecting *P* to the origin according to Eq. (4).

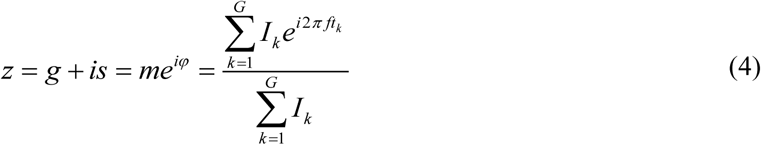

where *I*_*k*_ is the intensity of gate *k* (single pixel intensity for a single pixel phasor, or total intensity of the region of interest (ROI) for ROI analysis), and *f* is the phasor frequency. For a single-exponential decay, *P* is located on the universal semicircle (UC), and approaches (1, 0) when the lifetime tends to zero, and (0, 0) when the lifetime tends to infinity.

Fig. 3b illustrates the relation between the phase lifetime and the actual phasor *P* of the decay: the phase lifetime is the lifetime of the phasor located at the intersection of the UC and the line connecting *P* and the origin. The phase lifetime is obtained from the *g* and *s* coordinates of *P* as (19):

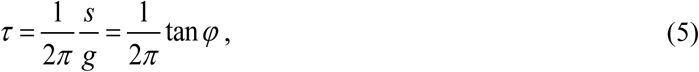

where *τ* is the lifetime, *s* and *g* are the phasor coordinates, and *f* is the phasor frequency. This expression has the advantage to be extremely simple to compute, and provides robust results even in the case of relatively low signal (28), in contrast to standard fitting methods. As discussed later, it also provides reliable estimates of the actual fluorescence lifetime for large range of acquisition parameters. In addition, since a phasor value can be defined for each pixel, it is possible to represent their individual phasors in a phasor plot and, by selecting specific regions in the phasor plot characterized by similar phasor values, highlight the location of the corresponding pixels in the original image, allowing a straightforward visual exploration of fluorescent lifetime heterogeneities within a sample.

#### Phasor calibration

In practice, the experimental gate shape and the time delay (offset) between the laser pulse and the trigger signal, both affect the shape of the decay recorded by the acquisition hardware, which is a convolution of the sample’s signal and the instrument response function (IRF). The effect of the IRF can be easily corrected using a calibration sample with a known lifetime *τ*_*cal*_. In the phasor representation, the presence of an IRF simply rescales the theoretical phasor’s modulus and rotates its phase by a fixed amount (the sample’s phasor is multiplied by the IRF’s phasor):

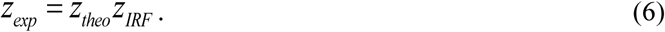

The IRF’s phasor can be obtained by measuring the calibration sample’s uncorrected phasor *z*_*cal,exp*_ and using the calibration sample’s theoretical phasor *z*_*cal,theo*_ (19):

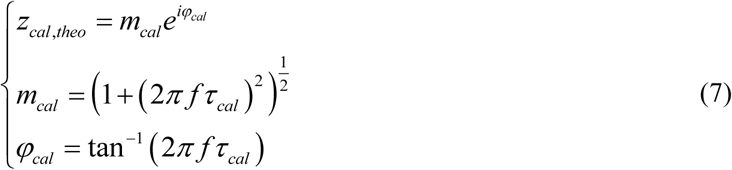

The same calibration approach in fact works generally well to correct for the decay modification brought about by the gating process, which amounts to an integration rather than a convolution:

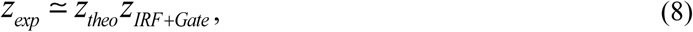

where *z*_*IRF+Gate*_ is a calibration factor incorporating both IRF and gate influences on the recorded decay. Eq. (8) worked satisfactorily in all cases studied in this work, and can be used provided the number of gates *G* is not too small (in practice, for *G* > 10) (23,27).

The calibration factor *z*_*IRF+Gate*_ can be calculated for each pixel (using a phasor calibration image), or for each ROI (using a phasor calibration “map”), or globally for the whole frame (single phasor calibration). In this study, we computed calibration factors for contiguous 4×4 pixel ROIs covering the whole field of view, due to the gate characteristics.

### 2.5. Photon Economy

An important figure-of-merit of a FLIM system is the optimality (or economy) with which the collected photons are used to extract the parameter of interest, namely the fluorescence lifetime *τ*. This can be quantified via the uncertainty *σ*_*τ*_ with which the lifetime is determined as a function of the number *N* of collected photons. While this parameter is insensitive to the speed of the system which determines the global photon count rate, it still quantifies its ability to operate in settings with scarce photons. A convenient measure of photon economy is the normalized relative error on the measured lifetime (29) or *F*-value (30) defined as:

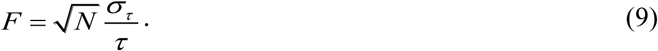

*F* = 1 in the case of a pure single-exponential decay measured using a TCSPC system without jitter and with a δ-function IRF (*i*.*e*. when the photon arrival times can be considered as the result of a pure Poisson process), and when the lifetime *τ* and its standard deviation *σ*_*τ*_ are calculated as the average of the *N* photon arrival times and their standard deviation. Departure from this value is expected when the measurement process is non-ideal (e.g. in the presence of jitter, finite IRF or detector/background noise) or when the lifetime is extracted using a different procedure (as is for instance the case in time-gated measurements, or using the phasor approach).

### 2.6. *F*-value dependence on gate width

As a result of the finite gate width *W*, the ideal *F*-value that can be achieved is no longer 1. The effect is analogous to what happens when departing from the ideal TCSPC case (for which *F* = 1) by addition of a photon timestamp uncertainty corresponding to binning (we neglect the uncertainty due to the finite IRF width and electronic jitter, whose contribution to the total variance does not play any significant role in the present experiments). An expression for *F* for the case of small gate width (appropriate for TCSPC methods with finite bin numbers or in for time-gating in the absence of gate overlap) was derived in (31) and used in later publications (e.g. (29,32,33)) but does not apply to the present situation involving large and overlapping gates. Pending a more rigorous analysis, a first order estimation of the effect of large gate width can be obtained from the following reasoning. Noting *W* the gate or bin width, and assuming, in a first approximation, that the photons collected in each gate are uniformly distributed, the resulting standard deviation of their timestamps, *σ*_*W*_, is given by (see supporting discussion for a detailed derivation):

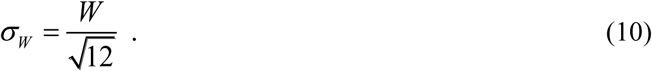

Similarly, the average of *N* such timestamps (which follow a Bates distribution) has a standard deviation given by:

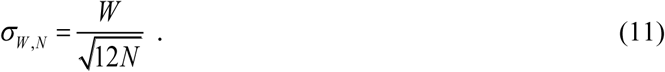

Since the time stamp uncertainty due to gating (or binning) is uncorrelated to the Poisson emission process, the variance of the two add up, resulting in a standard deviation of the lifetime, measured as an average of the binned timestamps, given by:

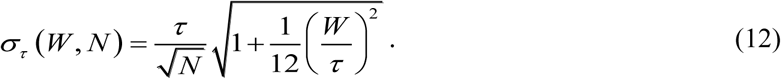

From the definition (9) of *F*, it results that:

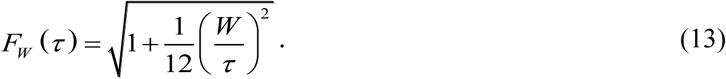

The above approximation is expected to become poorer as the gate width *W* becomes comparable to or larger than the measured lifetime, as the distribution of time stamps within a gate cannot be considered uniform anymore. However, Eq. (12) should provide a lower bound for the measured phase lifetime standard deviation.

In order to obtain a better estimation of the phase lifetime standard deviation and *F*-value expected in the case of such a gated counting process, we used Monte Carlo simulations of the emission and detection process, taking into account the gate number, gate width, laser period, lifetime, and total number of detected photons to compute the standard deviation of the phase lifetime obtained by phasor analysis. Because both calibration sample and sample of interest were independently acquired with a finite and generally different number of photons (*N*_*c*_ and *N*_*i*_, respectively), the phase lifetime variance of both samples were computed separately and summed to obtain the predicted standard deviation and *F*-value:

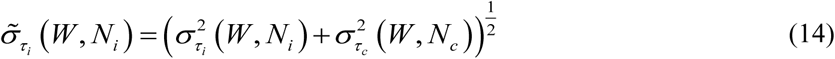

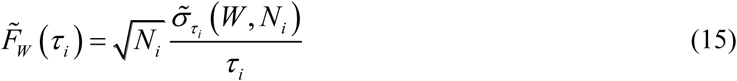

where *τ*_*c*_ and *τ*_*i*_ are the lifetimes of the calibration sample and sample of interest, respectively. This approach of summing the variance of the sample of interest and calibration sample was also used when comparing data and the shot noise model of Eq. (13).

### 2.7. Dye mixture analysis

The volume fraction in a mixture of two components characterized by different lifetimes, can be inferred from the location of its phasor in the phasor plot with respect to the pure components’ phasors, using the geometric phasor ratio (17,19). As illustrated in Fig. 4, the phasor of a mixture is located on the segment connecting the phasors of the two fluorescent dyes forming that mixture. The phasor fraction or ratio of dye 1 (*r*_*1*_) is expressed by the ratio of the distance between the phasor of the mixture and the phasor of the other dye (*d*_*2*_) to the total length of the segment (*d*_*1*_*+d*_*2*_):

**Fig. 4.**
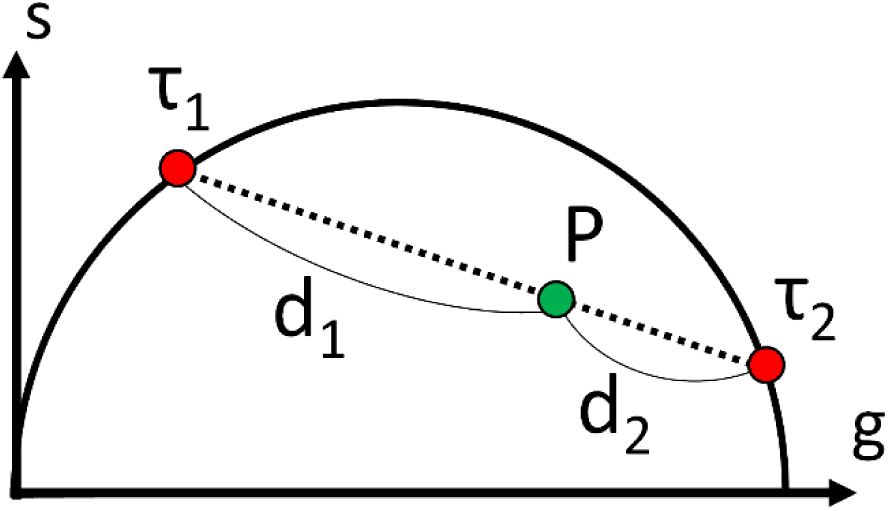
Conceptual illustration of mixture analysis. *P* is the phasor of the mixture, *τ*_*1*_ and *τ*_*2*_ are the phasors of two dyes, and *d*_*1*_ and *d*_*2*_ are the distances between the phasors of the dyes and the mixture. The phasor ratio can be found by calculating the ratio of the phasor distances, then can be converted to volume fraction using Eq. (16).

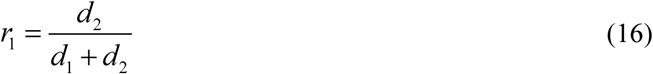

In dye mixture experiments, the phasors of all mixtures are first calculated as explained previously, followed by computation of the phasor ratio *r*_*1*_ of each ROI using Eq. (16) after projecting the phasors onto the segment connecting the pure samples’ phasors. The relation between the phasor ratio (*r*_*1*_) and the volume fraction (*v*_*1*_) is given by (23):

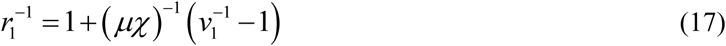

where *µ* is the initial concentration ratio and *χ* is the ratio of the product of extinction coefficient and quantum yield for each dye. The relation between *r*_*1*_ and *v*_*1*_ is therefore linear only if the product *µχ* = 1. In our experiment, in which a series of mixtures was prepared by using different volumes of two dye stock solutions, *μχ* is a constant, resulting in a simple relation between computed phasor ratio *r*_*1*_ and user-defined volume fraction *ν*_*1*_.

### 2.8. Data and software availability

All new raw data used in this manuscript is available in a public online repository at https://doi.org/10.6084/m9.figshare.8285993. This repository also includes data analysis files, Matlab scripts, and AlliGator analysis logs when relevant, as described in the Figshare Repository Description.pdf file. Software used in this work include Matlab (MathWorks, Natick, MA), Origin Pro 9.1 (OriginLab, Northampton, MA) and custom software developed in LabVIEW (National Instruments, Austin, TX). These programs are freely available as standalone Windows executables at:

- AlliGator (phasor analysis of time-gated data): https://sites.google.com/a/g.ucla.edu/alligator/
- Time-Gated Phasor Shot Noise Simulations: https://sites.google.com/a/g.ucla.edu/phasor-explorer/time-gated-phasor-shot-noise-simulations

## 3. Results

### 3.1. Time-gated data recording with SwissSPAD2

Fig. 5 shows the fluorescence decay profiles of four commercially available fluorescent samples used in the various experiments presented in this paper, as recorded by SS2 using a *W* = 13.1 ns gate width and 17.86 ps gate steps (total: 2,800 gates). ATTO 550, Cy3B and Rhodamine 6G (R6G) samples (Fig. 5a-c) were aqueous solutions sandwiched between two glass coverslips separated by a 1 mm thick rubber gasket, allowing to test the wide-field response uniformity of the detector. These samples were also used to study the dependence of phasor analysis performance on various acquisition parameters, as described in a later section. Fig. 5d shows the decay profile of an aqueous quantum dot (QD) sample (Qdot585 Streptavidin, ThermoFisher Scientific, ∼ 1 µM) left to dry out on a glass coverslip, resulting in random non-uniform density patterns, characterized by different average phase lifetimes across the field of view as discussed later. The three dye solutions had different concentrations (10 nM - 1 µM concentrations in aqueous buffer), ATTO 550 being the least concentrated, leading to noticeable shot noise, while this effect is minimal for the brighter sample, R6G. This variation provides an opportunity to investigate the effects of photon count on lifetime determination performance.

**Fig. 5.**
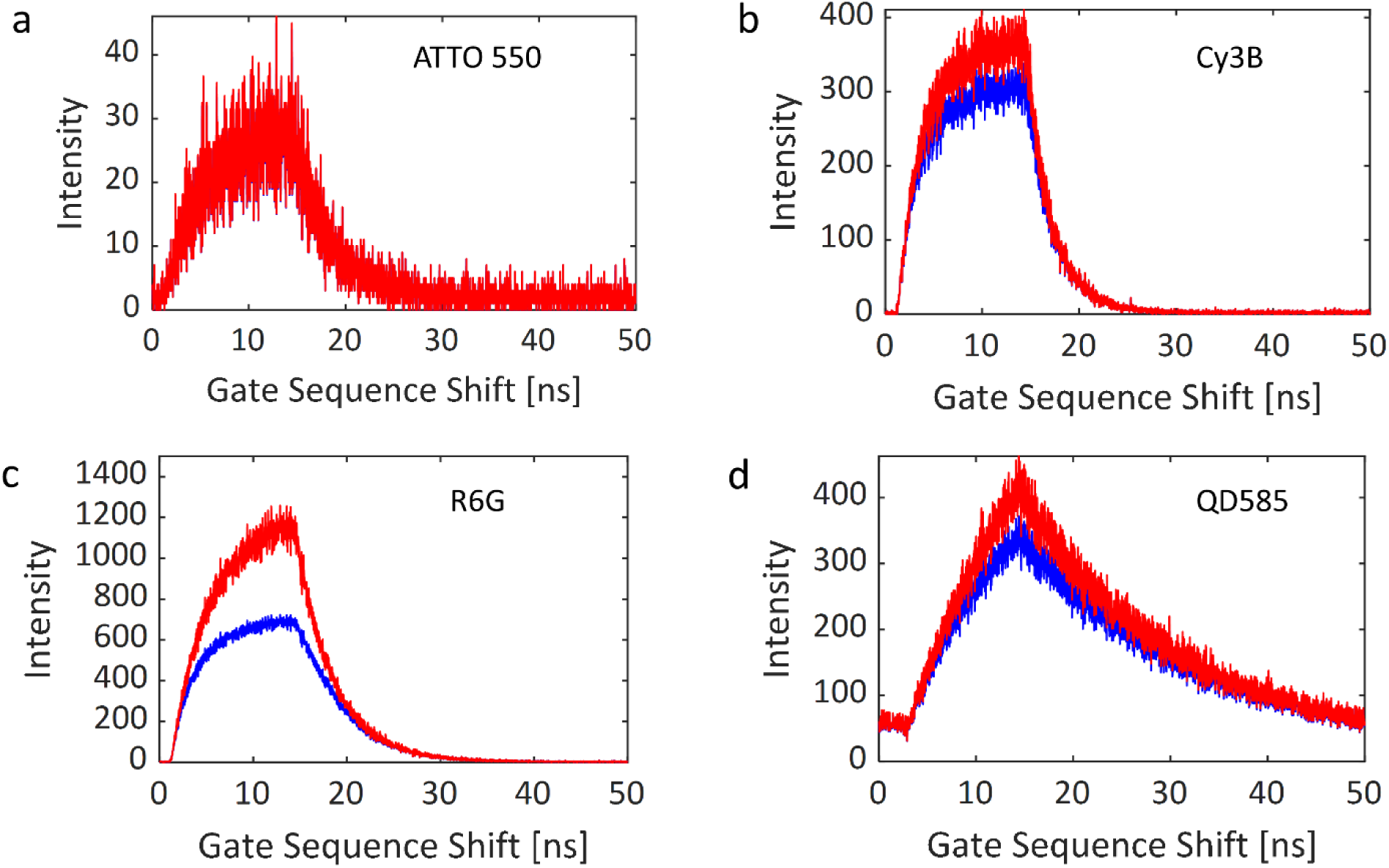
Gate intensity profiles (coordinates (193,190)) of (a) ATTO 550, (b) Cy3B, (c) Rhodamine 6G (R6G), and (d) quantum dot (QD585) solutions. Parameters: laser frequency: 20 MHz, gate width W = 13.1 ns, bit depth: 10, background correction: off. Blue: no pile-up correction, red: pile-up correction.

### 3.2. Phasor analysis of SwissSPAD2 data

The result of the phasor analysis of SwissSPAD2 data is illustrated in Fig. 6 with measurement of three fluorescent dyes with similar excitation and emission spectra (absorption peak around 550 nm, emission peak around 570 nm) but distinct lifetimes: Cy3B, R6G and ATTO 550 (literature values: *τ* = 2.8 ns, 4.08 ns and 3.6 ns respectively). In the experiment, all dye solutions were excited with a 532 nm 20 MHz pulsed laser characterized by ∼100 ps pulse width (LDH-P-FA-530XL, PicoQuant, Germany). ATTO 550 was selected as the intermediate lifetime species to compute a binned (4×4) calibration map used to calibrate the phasors of the two other dye samples (Cy3B & R6G) as explained in Methods. ATTO 550, Cy3B and R6G have similar photophysical properties but R6G is slightly brighter than the other two when excited at 532 nm (twice brighter than Cy3B and thrice brighter than ATTO 550). Moreover, because the concentration of the ATTO 550 sample was lower than that of the other two, we compensated its lower signal by a larger integration time (using 10-bit data instead of 8-bit data for the other two samples).

**Fig. 6.**
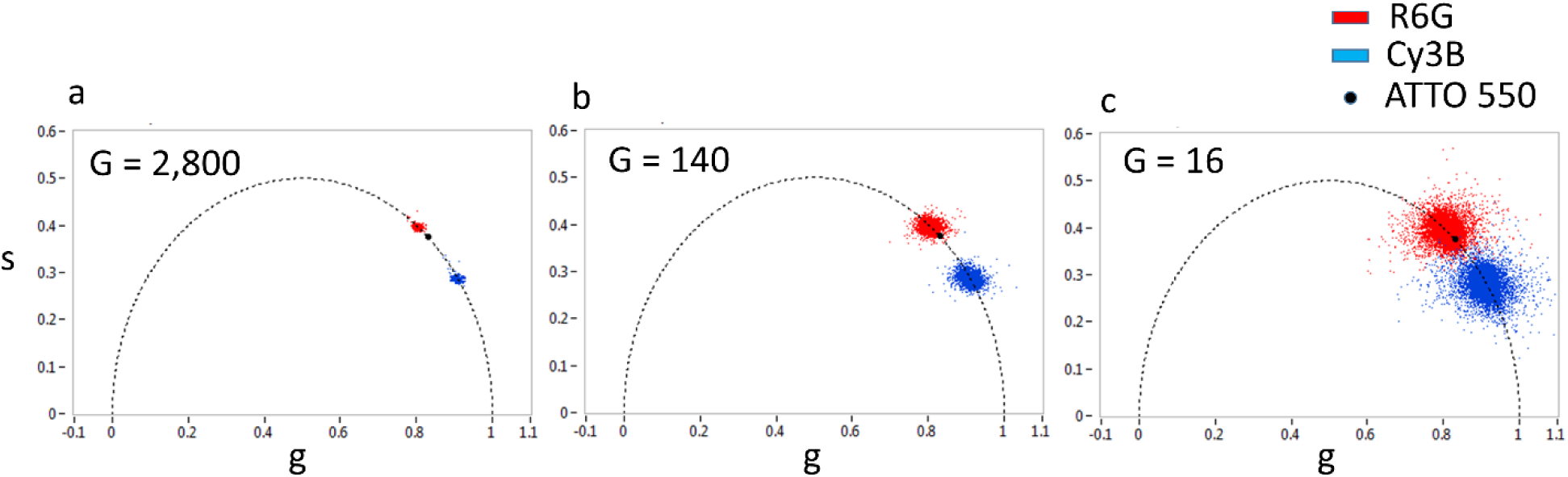
Phasor scatter plots for the R6G (τ = 4.08 ns) and Cy3B (τ = 2.8 ns) solutions obtained with 2,800 (a), 140 (b) and 16 (c) gate positions and calibrated with the corresponding ATTO 550 dataset (τ = 3.6 ns). The visual separation of the phasors of the two samples becomes more challenging when fewer gates (and thus fewer photons) are used. Even with as low as 16 gates, the two samples are clearly distinguishable. Experiment parameters: laser and phasor frequency: 20 MHz, gate width: 13.1 ns, array size: 472×256, binning: 4×4, bit depth: 8 (R6G & Cy3B), 16 (ATTO 550), pile-up correction: on, background correction: on, percentage of removed pixels: 0% (R6G & Cy3B), 0.5% (ATTO 550).

**Fig. 7.**
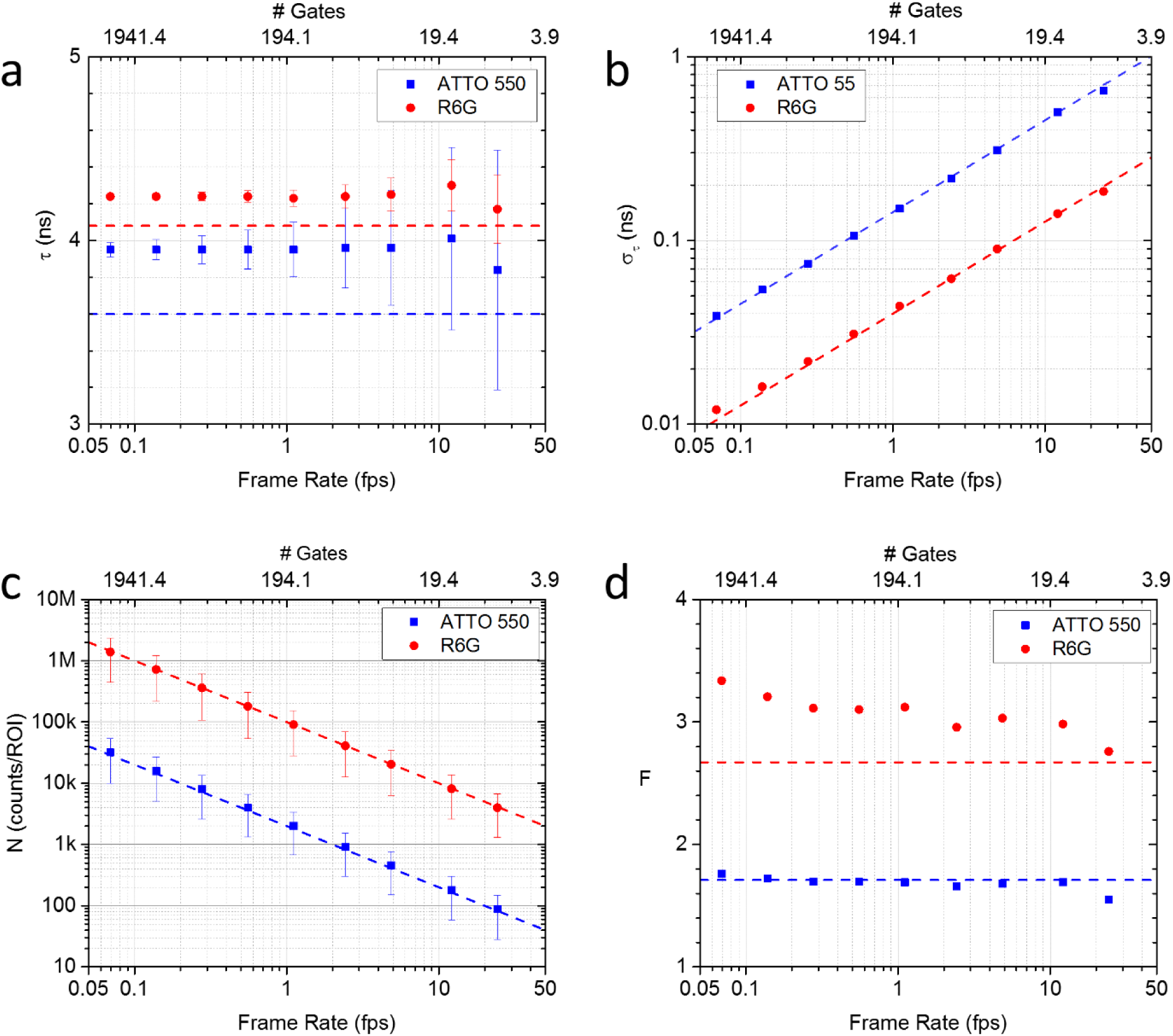
FLIM performance of SwissSPAD2 for different effective acquisition frame rates, determined by the num-ber of gate positions (# Gates = *G*). The numbers *G* used here are 2,800, 1,400, 700, 350, 175, 80, 40, 16 and 8: a: Phase lifetime ± standard deviation. The dashed lines indicate the literature values for both lifetimes. b: standard deviation. The dashed lines indicates a *G*^*-1/2*^ dependence. c: total photon counts per 4×4 pixel ROI. The dashed lines indicate a linear dependence on *G*. d: *F*-value of ATTO 550 and R6G data sets. The dashed lines indicate the Monte Carlo estimation of the effect of shot noise. Experimental parameters: laser & phasor frequency: 20 MHz, gate width: 13.1 ns, array size: 472×256, binning: 4×4, bit depth: 8, pile-up correction: on, background correction: on.

The phasor scatter plots calculated with 2,800, 140 and 16 gate positions are represented in Fig. 6a-c, respectively. The phasor dispersion clearly increases with decreasing gate number, due to the decreasing total signal. However, the two species are still resolvable with 16 gates, at an effective frame rate of 12.1 fps, showing that for these particular samples with a lifetime difference of 1.4 ns, the identification of dyes on the phasor map is possible at real-time acquisition speeds.

The phasor values can be converted into phase lifetime values (using Eq. (5)), resulting in normal distributions with mean and standard deviations provided in Table 2. The measured lifetimes show a slight negative bias (300 ps or 10% for Cy3B, 200 ps or 5% for R6G) compared to the literature values but matched those measured using a confocal TCSPC setup equipped with a different pulsed laser source (data not shown). Their standard deviation scales as *G*^*-1/2*^, where *G* is the number of gates used for the calculation, as expected for a shot noise-limited signal (Eq. (12)), since the number of counts is proportional to the number of gates used for analysis.

**TABLE 2:** Phase lifetime and standard deviation (in ns) obtained from Fig. 6. The measured phase lifetimes are slightly shorter than the literature values (Cy3B: 2.8 ns, R6G: 4.08 ns) and the standard deviation scales as G^-1/2^ as expected from Eq. (12).

| # Gates ($G$) | 2,800 | 140 | 16 |
| --- | --- | --- | --- |
| Cy3B | $2.49 \pm 0.02$ | $2.48 \pm 0.08$ | $2.43 \pm 0.23$ |
| R6G | $3.89 \pm 0.02$ | $3.88 \pm 0.08$ | $3.89 \pm 0.25$ |

### 3.3. Influence of frame rate on phase lifetime precision

To better characterize the performance of the phasor approach using SS2 data, we conducted a “virtual” experiment where frame rate increase was emulated by the reduction of the number of gates used to analyze the data. In principle, different acquisitions with decreasing number of gates could have been performed sequentially, but this would have exposed the sample to longer overall excitation, with possible detrimental effects such as bleaching. We investigated the corresponding effect of the number of gates on the accuracy and precision of phase lifetime extraction. However, because the number of recorded photons is proportional to the number of gates (at constant excitation intensity), this study is also a study of the dependence of these parameters on photon count. For this reason, we also measured the dependence of the *F*-value on the number of gates, as this parameter integrates the count dependency.

In this experiment, the lifetimes of ATTO 550 and R6G were measured with SwissSPAD2 using 2,800 gate positions. The original consecutive gates were separated by 17.86 ps and only one of the four 8-bit frames comprising each gate data was used (see Section 2.3). Each gate duration was 13.1 ns, encompassing approximately one quarter of the 50 ns laser period. Each binary frame was acquired using a 10 μs exposure (200 laser periods) followed by readout (10.2 μs) during which the detector was blind. This amounts to 8-bit gate image generation at 194 fps, or 14.42 s for a complete series of 2,800 gate images. The number of gate positions was then gradually reduced in post-processing by decimating gates from 2,800 down to 8, thus resulting in longer delay between the remaining consecutive gates. This process emulates raw data that would have been acquired faster if using longer gate delays and fewer gates. Phasors of all datasets were calibrated using the Cy3B dataset as reference.

The FLIM performance metrics for various numbers of gate positions are summarized in Fig. 7. Fig. 7a represents the average calibrated phase lifetime (Eq. (5)) of the 118×64 (4×4 pixel) ROIs plotted with their standard deviation as error bar. The literature lifetime values are represented as dashed lines for comparison. Up to an acquisition speed of 24.3 fps (*G* = 8 gates per decay), the lifetime was estimated with an accuracy better than 6.6 % for ATTO 550 and 2.2 % for R6G. Fig. 7b shows the phase lifetime standard deviation (precision), *σ*_*τ*_, exhibiting the expected *G*^-1/2^ scaling (or equivalently *N*^-1/2^, since the number of accumulated photons *N* is proportional to the number of gate *G* in these experiments, see Fig. 7c). The ability to achieve acceptable levels of precision down to 8 gates obviously depends on the total number of photons *N* in the data set. For this low number of gates, the relative error (*σ*_*τ*_/*τ*) is 4.5 % for R6G (the brighter sample), whereas it increases to 18.1 % for ATTO 550, which was recorded with approximately 45 times less photons, as shown in Fig. 7c. The relative error does therefore not scale exactly as expected from the signal ratio (3.9 vs v45 = 6.7), due to the additional effect of shot noise present in the calibration sample.

Fig. 7d shows the *F*-value of both samples as a function of effective frame rates. The *F*-value at 24.3 fps is 1.55 for ATTO 550 and 2.76 for R6G, respectively, and both remain approximately constant down to the lowest frame rate (or equivalently, up to the largest number of gates). These values compare well with the shot noise limits computed by Monte Carlo simulations (dashed lines), when the contribution of the calibration sample’s shot noise is included. The discrepancy is largest for the brighter R6G sample, for which the effect of shot noise in the calibration is most noticeable. These *F*-values express the fact that 1.55^2^ = 2.4 times more photons for ATTO 550 and 2.76^2^ = 7.6 times more photons for R6G must be detected compared to an ideal TCSPC FLIM system, in order to achieve the same lifetime precision. It is important to emphasize that these figures depend on the calibration sample used: because the Cy3B sample used in this analysis was dimmer than the R6G sample, a significant fraction of the *F*-value for R6G is due to the shot noise present in the calibration sample. Theoretically, with a very bright calibration sample (or a calibration sample measured using a long integration time), the *F*-value would scale as described by Eq. (13) and thus decrease as the measured lifetime increases.

Considering that these *F*-values are independent of frame rate (Fig. 7d), this result shows that time-gated phasor FLIM becomes competitive at high frame rates, which are challenging to achieve with scanning confocal TCSPC techniques.

### 3.4. Influence of time gate width on phase lifetime determination

Data required for the analysis of the influence of gate width *W* on lifetime was collected using seven distinct values of gate duration covering the achievable range of 10.8 ns to 22.8 ns, corresponding to approximately 1/4 to 1/2 of the laser period (*T* = 50 ns). The data was acquired with constant distance between successive gates (gate step) of 17.86 ps. This resulted in 2,800 gate positions over the laser period. Analyses were performed on two subsets of these 2,800 gates: one where 1 every 20 gates was retained (number of gates *G* = 140, separation: 357 ps), and the other in which 1 every 175 gates was used (*G* = 16, separation: 3.125 ns). The 472×256 pixel sensor was binned into adjacent 4×4 pixel ROIs to increase statistics. Cy3B (life-time: 2.8 ns) was used for phasor calibration.

The results of these analyses are shown in the 4 panels of Fig. 8. Fig. 8a shows that the extracted phase lifetime is independent from the gate width parameter *W* and exhibits a small positive bias compared to the literature values (ATTO 550: 3.94 ns vs 3.6 ns, R6G: 4.24 ns vs 4.08 ns).

**Fig. 8.**
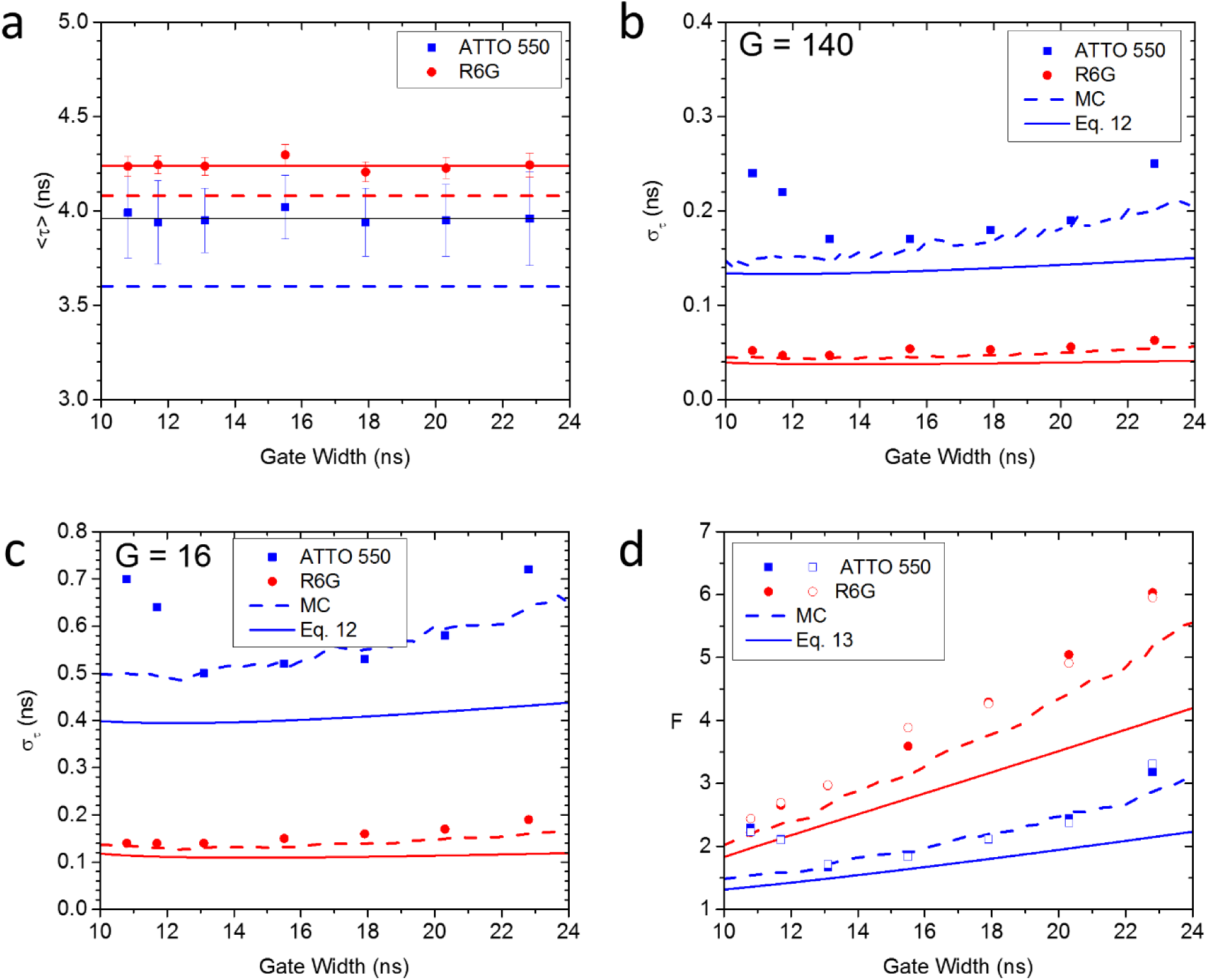
Dependence of the measured phase lifetime on gate width. a: Average phase lifetime of the ATTO 550 and R6G samples calibrated with the Cy3B sample (*τ* = 2.8 ns) using 140 gates. The points represent the average of all values in the image, while the error bars correspond to the measured standard deviation. The plain line correspond to the average of all values; the dashed line indicate the literature values for both dyes. b,c: Dependence of the phase lifetime standard deviation on gate width, for *G* = 140 (b) and *G* = 16 (c) gates. Points: measured values; plain lines: results of Eq. (13); dashed lines: MC results. d: Dependence of the F-value on gate width. Filled symbols: *G* = 16, open symbols: *G* = 140: plain lines: results of Eq. (14); dashed lines: MC results. Experimental parameters: laser & phasor frequency: 20 MHz, number of gate positions: 16 or 140, array size: 476×256, binning: 4×4, bit depth: 8, background correction: on, pile-up correction: on.

The standard deviation of the measured phase lifetime (Fig. 8b&c) depends on the number of gates used for the measurement (Fig. 8b: *G* = 140, Fig. 8c: *G* = 16) and increases slightly with the gate width above *W* = 13.1 ns. These trends are reproduced qualitatively by a simple shot noise model (Eq. (14), plain curves) of lifetime calculation by individual photon time stamp averaging, where the real time stamps are “blurred” by an amount equal to the gate width (see Methods). Because phase lifetime calculation involves a calibration step using a different sample characterized by different shot noise level, this simple model further assumes that their variances add up to yield the final measurement’s variance. While instructive, this model is clearly insufficient and will need to be refined further to better account for the actual gating process and analytical details of the phasor computation and calibration steps. In the meantime, we compared the measured standard deviation to simulated data taking into account the main characteristics of the measurement: exponential fluorescence decay with lifetime *τ*, finite number *G* of square time gates of width *W*, and finite number of recorded photons *N*. The results of these Monte Carlo calculations (performed on both sample of interest and calibration sample, followed by a summation of their variance) are indicated as dashed curves in Fig. 8b,c. While there is still some difference with the measured data, the agreement is satisfactory and suggests that this simple model captures the essential ingredients of our data acquisition and analysis process.

To separate the effect of shot noise from other effects, we looked at the *F*-value defined by Eq. (9). First, we verified that there was no residual dependence of the *F*-value on the number of photons *N*, by comparing the results obtained for the same data analysed with different number of gates (*G* = 140 & *G* = 16). Fig. 8d shows that there is no difference between measurements characterized by the same gate width but different gate numbers (open symbols: *G* = 140, plain symbols: *G* = 16), demonstrating that the contribution of *N* to the standard deviation is indeed of the form *N*^*-1/2*^. The remaining dependence is a monotonic increase with the gate width (except for a few anomalous values for small gate width in the case of ATTO 550). Indicating that at fixed detected number of photons N, it is preferable to use shorter gate to achieve a better precision. The simple shot noise model of Eq. (13) is indicated by plain curves, while the MC simulation results are represented as dashed curves. As for the standard deviation results of Fig. 8b,c, the shot noise model qualitatively reproduces the observation, but is fairly optimistic, whereas the numerical estimate is quantitatively correct.

### 3.5. Dye mixture analysis

Having demonstrated precise measurement of distinct lifetimes using SwissSPAD2, we examined its ability to quantify mixtures of two fluorescent dyes. While useful in and by itself in order to use lifetime as a contrast mechanism to distinguish between two species (instead of emission spectrum), this application is also relevant for some FLIM-FRET studies, in which the local fraction of FRET-undergoing donor molecules (characterized by a shorter lifetime than isolated donor molecules) is of interest(23). In the phasor method, the volume fraction of a species in the mixture is related to the phasor ratio, a quantity easily calculated as the relative distance of the mixture’s phasor to the pure species phasors (Eq. (17)). To validate this approach, a series of measurement of 5 mixtures of Cy3B and R6G solutions was performed (Fig. 9).

**Fig. 9.**
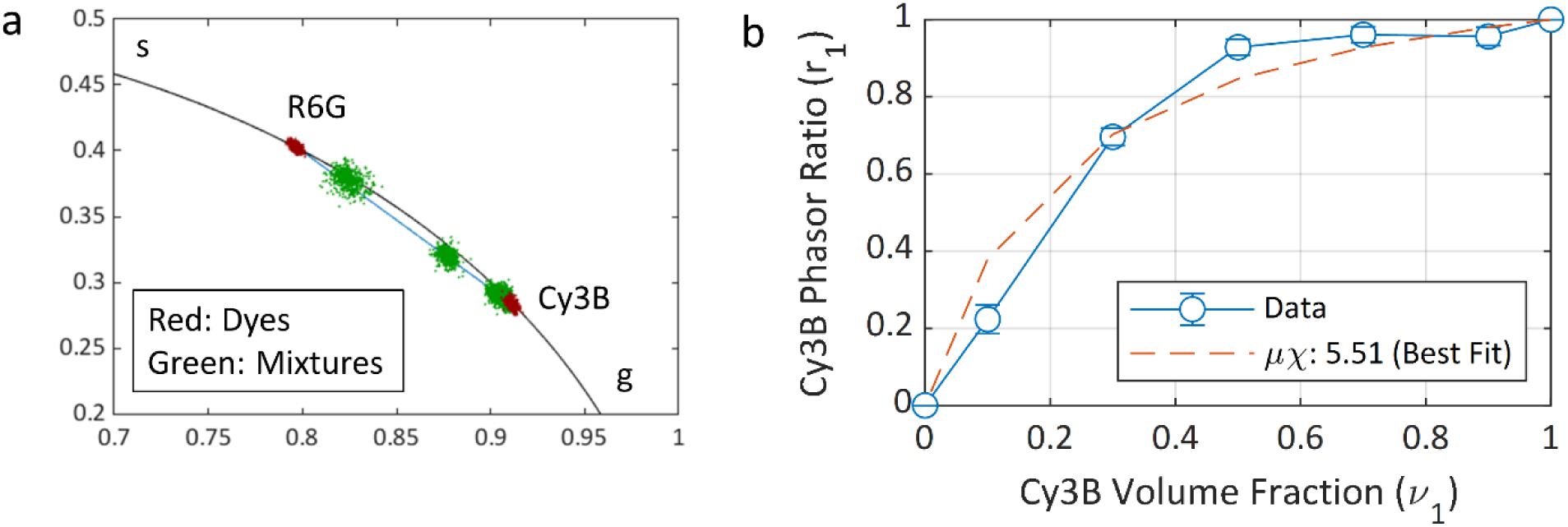
Dye mixture analysis of Cy3B and R6G with various volume fractions. A separate Cy3B sample was used as the reference dye for phasor calibration, using a *τ* = 2.5 ns (value measured by TCSPC). *μ* is the initial dye concentration ratio and *χ* is the product of the extinction coefficient ratio and quantum yield ratio for both dyes (23). (a) Phasors of the dyes (red) and mixtures (green) on the universal semicircle. (b) Calculated *μχ* for each mixture, and the *μχ* obtained by fitting method. Experimental parameters: laser PRF: 20 MHz, phasor frequency: 20 MHz, number of gate positions: 234, gate length: 22.8 ns, array size: 248×160, binning: 8×8, bit depth: 10, pile-up correction: on, background correction: on, percentage of removed pixels: 1%. Note that because the mixtures decays are not single-exponential, a constant background subtraction approach was used based on the background measured in the reference sample.

For this analysis, 248×160 of the brightest pixels were used, and 8×8 binning was performed to increase the signal-to-noise ratio. Fig. 9a shows the phasors of the dyes in red and the 5 mixtures in green, in which the Cy3B volume fraction varies from 10% to 90%. The higher initial concentration of Cy3B (∼ 1 µM) compared to that of R6G results in the phasors of the 50% to 90% Cy3B volume fractions to be virtually indistinguishable. Fig. 9b represents the phasor ratios extracted from Fig. 9a as a function of the Cy3B volume fraction. Fitting these data points to Eq. (17) yields *μχ* = 5.51 and *σ*_*μχ*_ = 1.77. Taking into account the fact that *χ* = 0.51 is the ratio of the product *ϕε* (*ϕ*: quantum yield, *ε*: extinction coefficient) for both dyes (23), this result shows that the initial concentration of the Cy3B solution is approximately *µ* = 10.8 times higher than that of R6G. This experiment therefore demonstrates the ability of phasor analysis of SwissSPAD2 data to perform quantitative mixture analysis.

### 3.6. Phase Lifetime Map for Complex Samples

A powerful aspect of the phasor method is its 2-dimensional representation of sample lifetimes in the phasor plot (Fig. 4). In this representation, species characterized by a single exponential decay with lifetime *τ* are all located in a well-defined region close to the universal circle (UC) and are easily distinguishable from samples with different lifetimes (see for instance Fig. 6). For a sample comprising two fluorescent species with different lifetimes localized in separate regions of the image, the phasor plot will exhibit two separate clusters of phasors in the phasor plot. It is then simple to correlate the location of phasors in the phasor plot and the position of the corresponding source within the image, for instance by color-coding pixels whose phasors are in the first region of the phasor plot, say, red, and pixels whose phasor are in the other, say, green. In any other situation (for instance when different species with distinct lifetimes are present but colocalized in the image), the phasors will be intermediate (see Fig. 9) and interpretation in terms of lifetime is more subtle, but a similar and useful color-mapping can still be used, as illustrated next.

To illustrate this practical aspect of the phasor plot in the particular case of the SwissSPAD2 sensor, we studied a sample of commercial quantum dots emitting in the same spectral range as the organic dye samples discussed previously (Qdot 585 Streptavidin, peak emission wavelength: 585 nm), but with much longer lifetimes (see Fig. 5d). In addition to be characterized by longer lifetimes, quantum dots (QDs) generally exhibit size polydispersity, which results in heterogeneous photophysical properties (such as emission spectrum peak, lifetime, etc.) (34), which also depend on their environment.

10 µL of a stock solution of this QD sample (concentration ∼ 1 µM) was left to dry on a coverslip and imaged in ambient conditions using the same setup as used in the previous measurements. The corresponding intensity image is shown in Fig. 10a, which is characterized by bright random stripes of high QD concentration, interspersed with regions (stripes and dried out microdroplet regions) of lower concentration. The phasors of each pixel in this image (calibrated with the Cy3B sample described in the previous sections) are represented as a 2-dimensional histogram shown in Fig. 10d. While there is a noticeable dispersion in the observed phasors, they are close to the UC, and aligned along a straight line between two phasors (indicated by a red and green dot respectively) with phase lifetime *τ*_*R*_ = 13.9 ns and *τ*_*G*_ = 16.7 ns. Using the relative distance of each phasor to these two references (or phasor ratio *r*_*G*_, Eq. (16)) (19) to color-code the original pixels in the source image (*r*_*G*_ = 0: red, *r*_*G*_ = 1: blue, spectrum color scale in between, as indicated in Fig, 10b) yields the phasor map shown in Fig. 10b, where the color of each pixel corresponds to its phasor ratio.

**Fig. 10.**
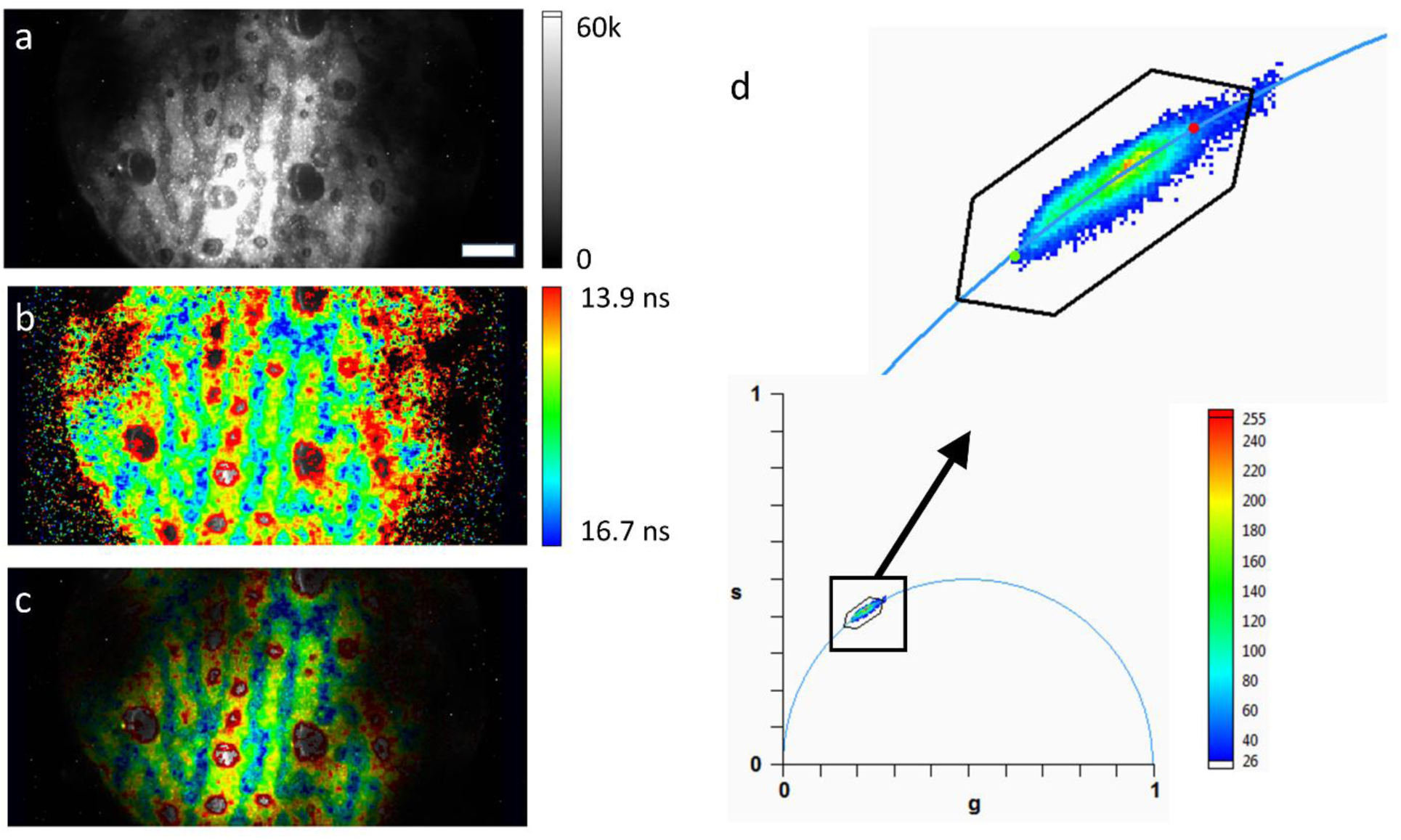
QD phase lifetime map. a: Intensity image of a dried QD sample. The contrast has been adjusted to be able to see most of the field of view. Scale bar: 25 μm. b, c: Color-coded phase lifetime maps. Two references (green dot: 16.7 ns and red dot: 13.9 ns) were defined in the phasor plot shown in d. Pixels were color-coded according to the their phasor ratio with respect to these two references and using the “spectrum” color scale indicated in b. Pixels with phasors close to the first reference (green dot: longer lifetimes) were colored blue, while pixels with phasors close to the second reference (red dot: shorter lifetimes) were colored red. Pixels with phasors in between were colored with an intermediate color. Points outside the segment were colored according to the closest point on the segment. The elongated hexagon represents the boundary of the region of the phasor plot were this color coding scheme applies. In b, the luminance is kept identical for all pixels, irrespective of their actual intensity allowing to visualize low intensity pixels (and their phase lifetimes). There is no obvious correlation between lifetime and intensity, while there appears to be a correlation between concentration and lifetime. In c, the luminance of each pixels scales with their intensity (shown in a). d: Bottom: Phasor plot of the data shown in a. Top: detail of the square region selected in the bottom phasor plot. The two references (green and red dots) are visible at both extremities of the phasor cloud.

Close comparison of Fig. 10a and Fig. 10b shows that dim regions at the periphery of the field of view are of no particular color and are associated with either long or short lifetimes, excluding a direct correlation between excitation intensity and observed lifetime, and thus, the possibility that the observed differences are due to imperfect background subtraction. On the other hand, bright stripes in Fig. 10a (corresponding to more concentrated regions of the sample), appear to be associated with shorter lifetimes (red/yellow color in Fig. 10b) than intermediate regions (blue/green, longer lifetimes, corresponding to lower QD concentrations). Fig. 10c combines the information provided by the other two, using the intensity image to scale the luminance of the color-coded phasor map, a conventional representation in standard FLIM imaging approaches.

## 4. Discussion

In this work, we examined the performance of a new wide-field time-gated SPAD array for FLIM using the phasor approach. In this sensor, we implemented relatively long time gates (11,12,20). This design choice, necessary to reduce the pitch of each pixel and to scale up the array to a large format, departs from most time-gated detectors, which strive to achieve the shortest possible gate duration to emulate the performance of TCSPC techniques. Moreover, the data content of each pixel is 1 bit, corresponding to one or zero photon count per readout period. This property of the sensor results in the need for pile-up correction at the pixel level.

This study demonstrates that time-gated imaging with very long time-gates, i.e. several times the typical lifetime of most organic fluorophores, and in regimes where pile-up effects are significant, does not preclude precise determination of fluorescence lifetimes, provided sufficient signal is recorded and gate boundaries are well defined. We obtained these results using a simple approach to phasor analysis, where calibration is performed using by a sample with known lifetime by a single algebraic operation (Eq. (8)), which is adequate down to a surprisingly small number of gates (*G* = 8 as tested in this work, see Fig. 7), at the expense of a minimal bias of the measured phase lifetime (160-400 ps, Fig. 7&8).

Examining the quantitative dependence of the measured lifetime’s standard deviation on the different user-selectable parameters: gate width *W*, gate number *G* and total signal *N*, we showed that this approach is essentially shot noise-limited. A simple analytical model assuming independent contributions from photon arrival time averaging and blurring due to the gate width (Eq. (12), Fig. 8) provides a lower bound to the measured lifetime’s standard deviation. A better and in some cases excellent estimate is provided by numerical simulation of the cumulated effects of shot noise (finite number of detected photons) and phasor calibration by a shot noise-limited sample (Eq. (14), Fig. 8). Residual effects of detector jitter non-uniformity, additional uncertainty due to pile-up correction and other possible factors related to phasor calibration could potentially further improve the agreement between observed and predicted lifetime uncertainty.

The overall performance of a wide-field time-resolved imaging system can be defined in two different ways. The first way is to find the required time to determine a lifetime with a given precision for a fixed pixel area, with no restrictions on the illumination level. This parameter is determined by the reduced photon economy (expressed by the normalized relative error on the measured lifetime or *F*-value, Eq. (9)) and the maximum local photon count rate. In some cases, however, there is a limit on the acceptable illumination level. Provided that the emission intensity is within the dynamic range of the detector, the performance of the system in this case is determined by the sensitivity of the pixel and the photon efficiency. To define the current and potential capabilities of SwissSPAD2, the limitations for each of these parameters must be well understood.

By definition, maximum local count rate is the inverse of the minimum delay between two detectable photons. In SwissSPAD2 operating in global shutter mode, this delay is equal to the sum of exposure and readout times, since the in-pixel memory can only store a single photon. If more than one photon is detected by the SPAD in a single frame, all photons except the first one are missed. This phenomenon, called pile-up, leads to distortions in the fluorescence decay shape. The pile-up correction used here (Eq. (1)) partially recovers the photon distribution; however they cannot improve the signal-to-noise ratio degradation caused by missed photons. To minimize pile-up, the average photon count per binary frame must be kept significantly below 1 for all gates. To find the maximum local count rate, the maximum allowed photon count for acceptable pile-up must be multiplied by the ratio between the average and peak intensity of the gate response, which is influenced by the sample lifetime, gate length and laser pulse width. Increasing the bit depth of the in-pixel memory or the readout speed are two possible ways of increasing the maximum count rate, at the expense of increased afterpulsing (which would show up as background noise for FLIM purposes). Both solutions demand additional chip area, therefore they come with losses in fill-factor or increase in pixel size.

The sensitivity of the imager is determined by the SPAD photon detection efficiency (PDE) and the pixel dead time. The PDE is the percentage of incoming photons that are detected by the SPAD, which is equal to the product of the PDP and fill factor. To improve the PDE, microlenses were deposited on the imager. The concentration factor of the microlenses on SwissSPAD2 was measured between 2.6 (*V*_*ex*_ = 6.5 V) and 4.5 (*V*_*ex*_ = 3.5 V), resulting in an effective fill-factor between 28% and 47%. The dead time is the time during the sensor operation where it is not sensitive to photons. In SwissSPAD2 operating in low pile-up regime, the dead time consists of the readout time if it is in the global shutter mode, and the duration of the laser period where the gate is closed. The first problem can be solved by switching to rolling shutter mode where the exposure and readout occur simultaneously. For the current version of the chip, this operation requires improvements in power distribution network. The second problem can be solved by adding a second gate to the pixel (35). This dual-gate architecture enables the pixel to be sensitive during the entire laser period, while recording timing information by assigning the photon to one of the two gates. With these two additions, the dead time can be virtually eliminated.

## 5. Conclusion

The capability of achieving good lifetime accuracy and high precision (140 ps) at close to video rate (12.4 fps, Fig. 7b) is an encouraging milestone towards real-time FLIM. In order to fully achieve this target, the next step will involve on-FPGA implementation of phasor calculation, enabling high-throughput data processing. The extraction of species fraction in complex mixtures (Fig. 9) is also an important step towards applications to FLIM-FRET measurements. The combination of improved sensitivity, real-time time-gated imaging and multiple species quantification will extend the capabilities of this sensor to applications demanding both high speed and high precision, such as small animal imaging.

## 6. Acknowledgments

This work was supported in part by the Swiss National Science Foundation Grant 166289 and the Netherlands Organization for Scientific Research Project 13916 (EPFL), and by HFSP Grant RGP0061/2015, NIH Grant GM 095904 and CRCC Grant CRR-18-523872 (UCLA). The authors have declared that no conflicting interests exist.

