## Supporting Information for "Wide-field time-gated SPAD imager for phasor-based FLIM applications"

Content:
- Supplementary discussion
- Supplementary table
- Supplementary figures
- Supplementary references

### SUPPLEMENTARY DISCUSSION

In this supplement, we provide the derivation of Eq. (11)-(13) in the main text.

Consider arrival times $\{t_i\}$ uniformly distributed in an interval [0, *W*].

Their average $<t_i>$ is given by:

$$\langle t_i \rangle = \frac{1}{W}\int_0^W t dt = \frac{1}{W}\left[\frac{t^2}{2}\right]_0^W = \frac{W}{2} \quad (1)$$

Similarly, the average of their squares is given by:

$$\langle t_i^2 \rangle = \frac{1}{W}\int_0^W t^2 dt = \frac{1}{W}\left[\frac{t^3}{3}\right]_0^W = \frac{W^2}{3} \quad (2)$$

Their variance is therefore given by:

$$\langle t_i^2 \rangle - \langle t_i \rangle^2 = \frac{W^2}{12} \quad (3)$$

which yields Eq. (10) of the main text.

We next compute the average of *N* such uniformly distributed arrival times:

$$\theta_N = \frac{1}{N}\sum_{i=1}^{N} t_i \quad (4)$$

The random variable (RV) $\theta_N$ defined as above is distributed according to the Bates distribution (1):

$$p(\theta; N) = \frac{N}{2(N-1)!}\sum_{k=0}^{N}(-1)^k \frac{N!}{k!(N-k)!}(N\theta - k)^{N-1}\sigma(N\theta - k) \quad (5)$$

where $\sigma(x)$ is the sign function ($\sigma(x)$ = -1 if $x$<0, $\sigma(x)$ = 1 if $x$>0, $\sigma(0)$ = 0).

The mean and variance of the Bates distribution are given by:

$$\begin{cases} \langle \theta_N \rangle = \dfrac{W}{2} \\ \langle \theta_N^2 \rangle - \langle \theta_N \rangle^2 = \dfrac{W^2}{12N} \end{cases} \quad (6)$$

from which Eq. (11) of the main text follows.

In the case of photons corresponding to an exponentially decaying fluorescent source with lifetime $\tau$, the arrival times $\{t_i\}$ detected by a gated detector with gate width W and gate boundaries [$\mu_j$ -$W$/2, $\mu_j$ + $W$/2] can be decomposed into:

$$\begin{cases} t_i = \mu_{j_i} + \delta t_i \\ \delta t_i \in \left[ -\frac{W}{2}, \frac{W}{2} \right] \end{cases} \tag{7}$$

where the gate center locations, $\mu_j$, are exponentially distributed with mean and variance $\tau$, and the $\delta\tau_i$ are uniformly distributed. $\theta_N$, the RV defined above as the average of $N$ such arrival times, can thus be written as the sum of two independent RVs:

$$\theta_N = \frac{1}{N}\sum_{i=1}^{N} \mu_{j_i} + \frac{1}{N}\sum_{i=1}^{N} \delta t_i \tag{8}$$

Because these two random variables are independent, their mean and variance add up:

$$\begin{cases} \langle \theta_N \rangle = \tau \\ \langle \theta_N^2 \rangle - \langle \theta_N \rangle^2 = \frac{\tau^2}{N} + \frac{W^2}{12N} \end{cases} \tag{9}$$

To obtain the above result, from which Eq. (13) of the main text follows, we have used the fact the first RV in Eq. (8) is the average of $N$ exponentially distributed RV. The sum of $N$ identical exponentially distributed RVs is distributed according to an Erlang distribution (2):

$$Erl(x; N, \tau) = \frac{}{(N-1)!\tau} \left( \frac{x}{\tau} \right)^{N-1} \exp\left( -\frac{x}{\tau} \right) \tag{10}$$

Its mean and variance are given by:

$$\begin{cases} \langle Erl(x; N, \tau) \rangle = N\tau \\ \mathrm{var}(Erl(x; N, \tau)) = N\tau^2 \end{cases} \tag{11}$$

from which the mean and variance of $1/N$ the same RV (which intervenes in Eq. (8)) follow.

### SUPPLEMENTARY TABLE

**TABLE S1:** Parameters of 7 gate configurations tested for time-resolved imaging firmware of SwissSPAD2. The gate length *W* characterizes the exposure time per laser period ($T$ = 50 ns in these experiments) and may differ slightly across the array (dispersion characterized by a full-width-at-half-maximum – FWHM) reported in ps. The theoretical square profile of the gate is in practice replaced by a plateau preceded by a finite rise time and followed by a fall time, both of which vary slightly across the array. This dispersion is characterized by a mean rise and fall times, as well as a rise and fall time FWHM reported in ps.

| | **Gate Configuration** | | | | | | |
|---|---|---|---|---|---|---|---|
| | **1** | **2** | **3** | **4** | **5** | **6** | **7** |
| **Gate Length (mean) [ns]** | 10.8 | 11.7 | 13.1 | 15.5 | 17.9 | 20.3 | 22.7 |
| **Gate Length (FWHM) [ps]** | 306 | 256 | 226 | 166 | 139 | 120 | 112 |
| **Rise Time (mean) [ps]** | 320 | 383 | 378 | 333 | 371 | 339 | 375 |
| **Fall Time (mean) [ps]** | 618 | 616 | 617 | 617 | 622 | 617 | 615 |
| **Rise Time (FWHM) [ps]** | 60 | 65 | 70 | 64 | 56 | 57 | 62 |
| **Fall Time (FWHM) [ps]** | 80 | 79 | 79 | 79 | 79 | 79 | 79 |
| **Rising Edge Position (FWHM) [ps]** | 319 | 269 | 252 | 193 | 185 | 161 | 148 |
| **Falling Edge Position (FWHM) [ps]** | 219 | 219 | 221 | 216 | 218 | 220 | 218 |

### SUPPLEMENTARY FIGURES

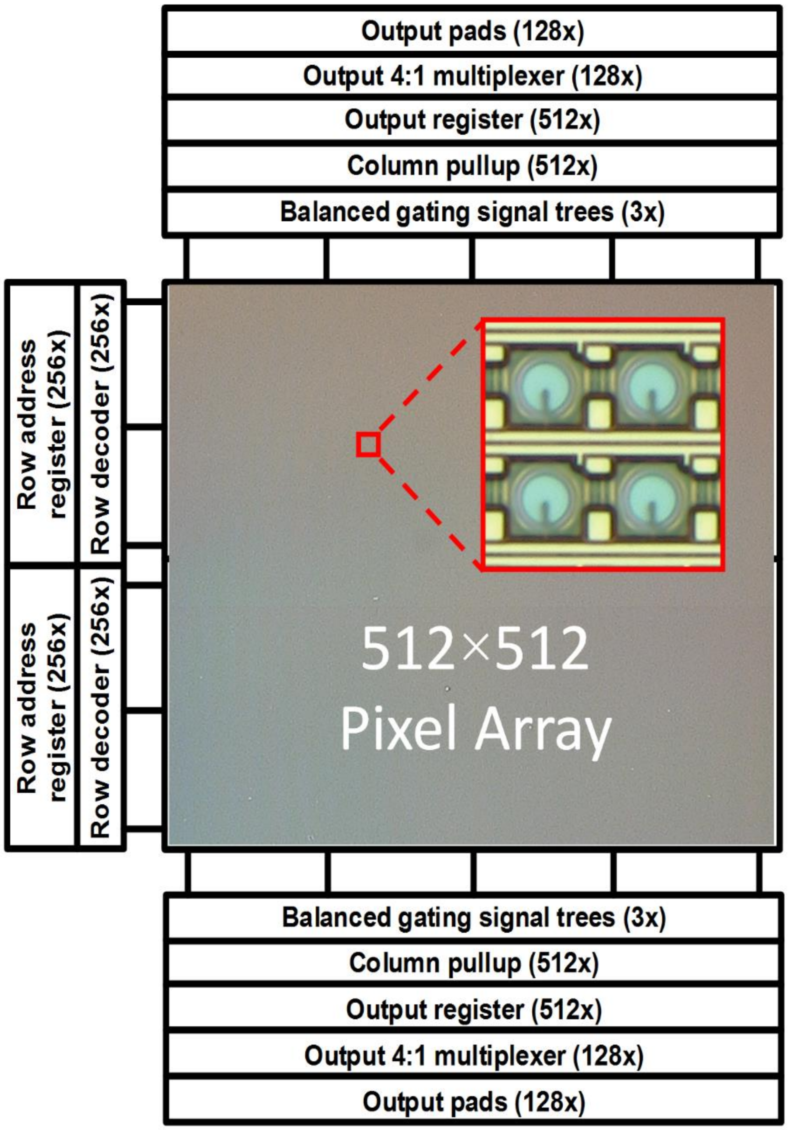


**Fig. S1.** Die micrograph and block diagram of the SwissSPAD2 image sensor (3).

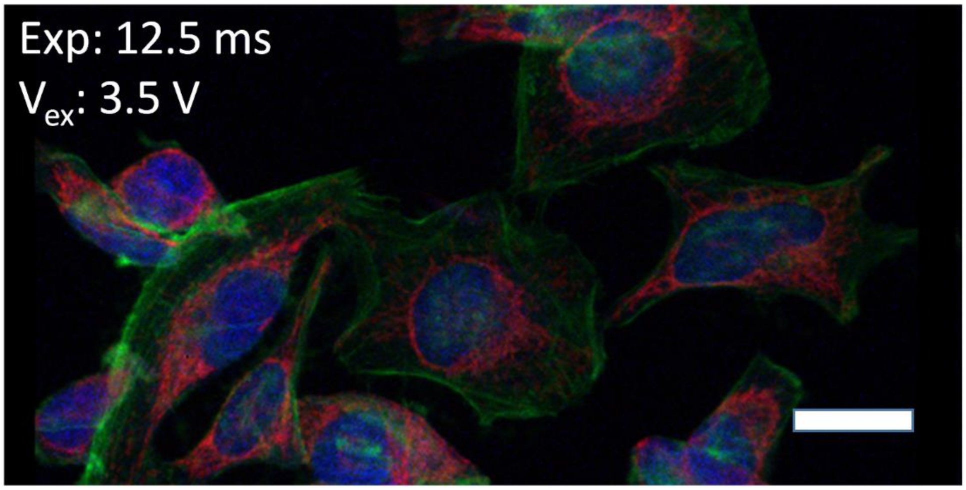


**Fig. S2.** A multicolor fluorescence intensity image of HeLa cells labeled with 3 different fluorescent dyes, captured by SwissSPAD2 (3). Scale bar is 26.2 μm.

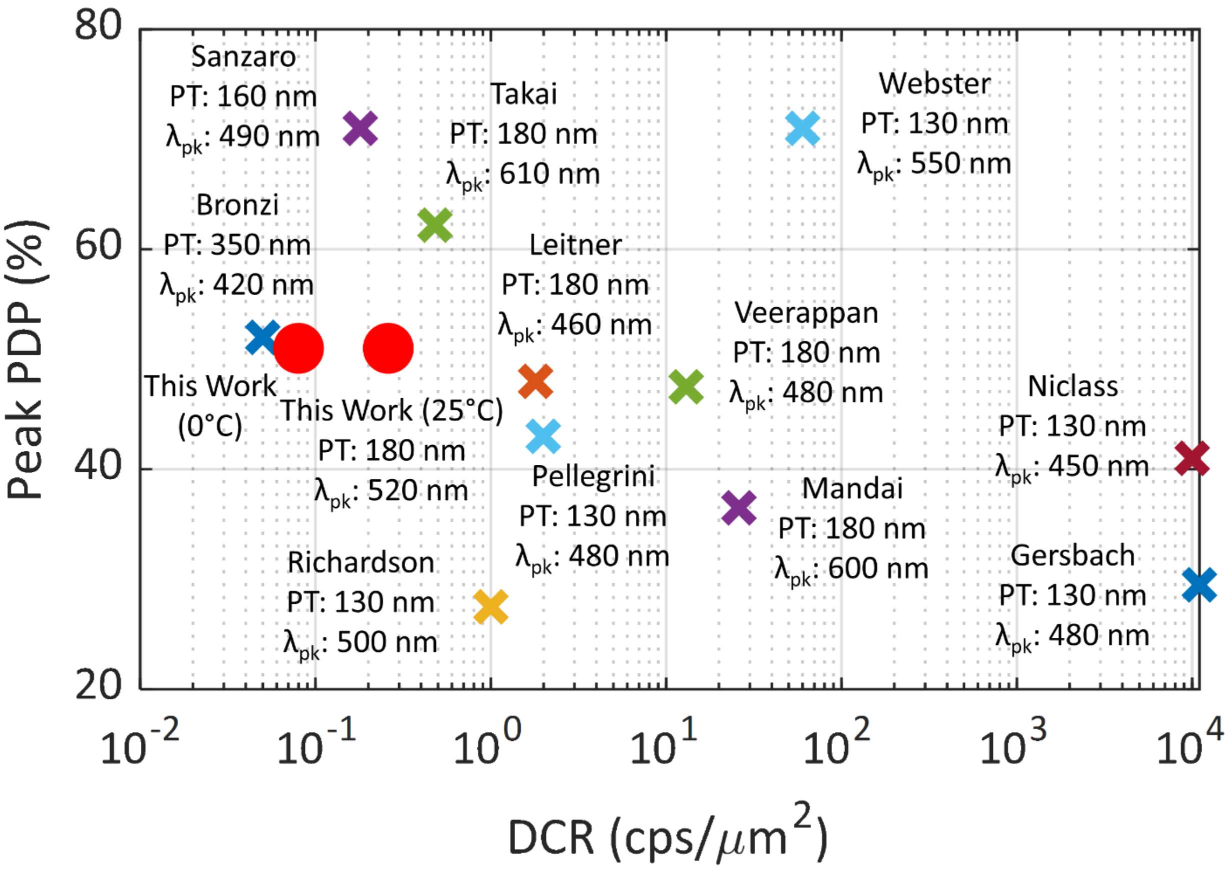


**Fig. S3.** PDP and DCR performance of SPADs fabricated in standard CMOS process technologies (3). (PT: Process technology, $\lambda_{pk}$: Peak wavelength). References cited : (4-14).

Sample 1

| | | | | |
|---|---|---|---|---|
| -0.001 | 0.007 | 0.007 | 0.006 | -0.001 |
| 0.004 | 0.034 | 0.057 | 0.032 | -0.003 |
| 0.000 | 0.056 | 100 | 0.059 | 0.001 |
| 0.001 | 0.030 | 0.053 | 0.029 | 0.003 |
| -0.001 | 0.006 | 0.004 | 0.000 | 0.001 |

Sample 2

| | | | | |
|---|---|---|---|---|
| 0.004 | 0.004 | 0.003 | 0.001 | 0.002 |
| -0.001 | 0.030 | 0.059 | 0.031 | 0.007 |
| 0.012 | 0.056 | 100 | 0.073 | 0.001 |
| 0.007 | 0.032 | 0.056 | 0.031 | 0.003 |
| -0.005 | 0.002 | 0.008 | -0.001 | 0.000 |

Sample 3

| | | | | |
|---|---|---|---|---|
| 0.002 | 0.000 | 0.006 | 0.000 | 0.002 |
| 0.005 | 0.028 | 0.054 | 0.029 | 0.005 |
| 0.003 | 0.061 | 100 | 0.064 | 0.009 |
| 0.000 | 0.032 | 0.071 | 0.032 | -0.002 |
| 0.005 | 0.001 | 0.008 | -0.003 | 0.002 |

Sample 4

| | | | | |
|---|---|---|---|---|
| -0.003 | 0.006 | 0.006 | -0.001 | 0.000 |
| -0.003 | 0.026 | 0.056 | 0.032 | 0.004 |
| 0.007 | 0.060 | 100 | 0.055 | 0.007 |
| -0.001 | 0.026 | 0.055 | 0.029 | 0.002 |
| -0.003 | 0.004 | 0.003 | 0.001 | -0.004 |

Sample 5

| | | | | |
|---|---|---|---|---|
| 0.006 | 0.001 | 0.005 | 0.006 | -0.005 |
| 0.005 | 0.033 | 0.062 | 0.032 | 0.005 |
| 0.013 | 0.067 | 100 | 0.065 | 0.008 |
| 0.000 | 0.035 | 0.066 | 0.025 | -0.002 |
| 0.009 | 0.003 | 0.003 | 0.002 | 0.003 |

**Fig. S4.** Crosstalk characterization results of five SwissSPAD2 samples at 6.5 V excess bias voltage. In each matrix, the values indicate the crosstalk probability of the pixels surrounding the reference pixel in the center. The values are calculated as the normalized median photon counts of multiple hot pixels and their neighbors in a dark image. The median dark count was subtracted from all pixels before the calculation. The results show that the crosstalk probability between nearest neighbors is less than 0.075%.
